# *hoxb* genes determine the timing of cell ingression by regulating cell surface fluctuations during zebrafish gastrulation

**DOI:** 10.1101/2024.02.06.579056

**Authors:** Yuuta Moriyama, Toshiyuki Mitsui, Carl-Philipp Heisenberg

## Abstract

During embryonic development, cell behaviors need to be tightly regulated in time and space. Yet, how the temporal and spatial regulations of cell behaviors are interconnected during embryonic development remains elusive. To address this, we turned to zebrafish gastrulation, the process where dynamic cell behaviors generate the three principal germ layers of the early embryo. Here, we show that *hoxb* cluster genes are expressed in a temporally collinear manner at the blastoderm margin of the embryo to regulate the timing of mesoderm and endoderm (mesendoderm) cell ingression. Under- or over-expression of *hoxb* genes perturb the timing of mesendoderm cell ingression and, consequently, the positioning of these cells along the forming anterior-posterior body axis. Finally, we found that *hoxb* genes control the timing of mesendoderm ingression by regulating cell surface fluctuations. Thus, *hoxb* genes interconnect the temporal and spatial pattern of cell behavior in the early embryo by controlling cell surface fluctuations.

**Highlights:**

- *hoxb* gene expression shows temporal collinearity at the blastoderm margin during zebrafish gastrulation.
- Temporal collinear expression of *hoxb* genes at the blastoderm margin regulates the timing of cell ingression and delineates spatial collinearity after gastrulation.
- *hoxb* genes regulate cell surface fluctuations and bleb formation at the blastoderm margin during cell ingression

**Graphical abstract:** 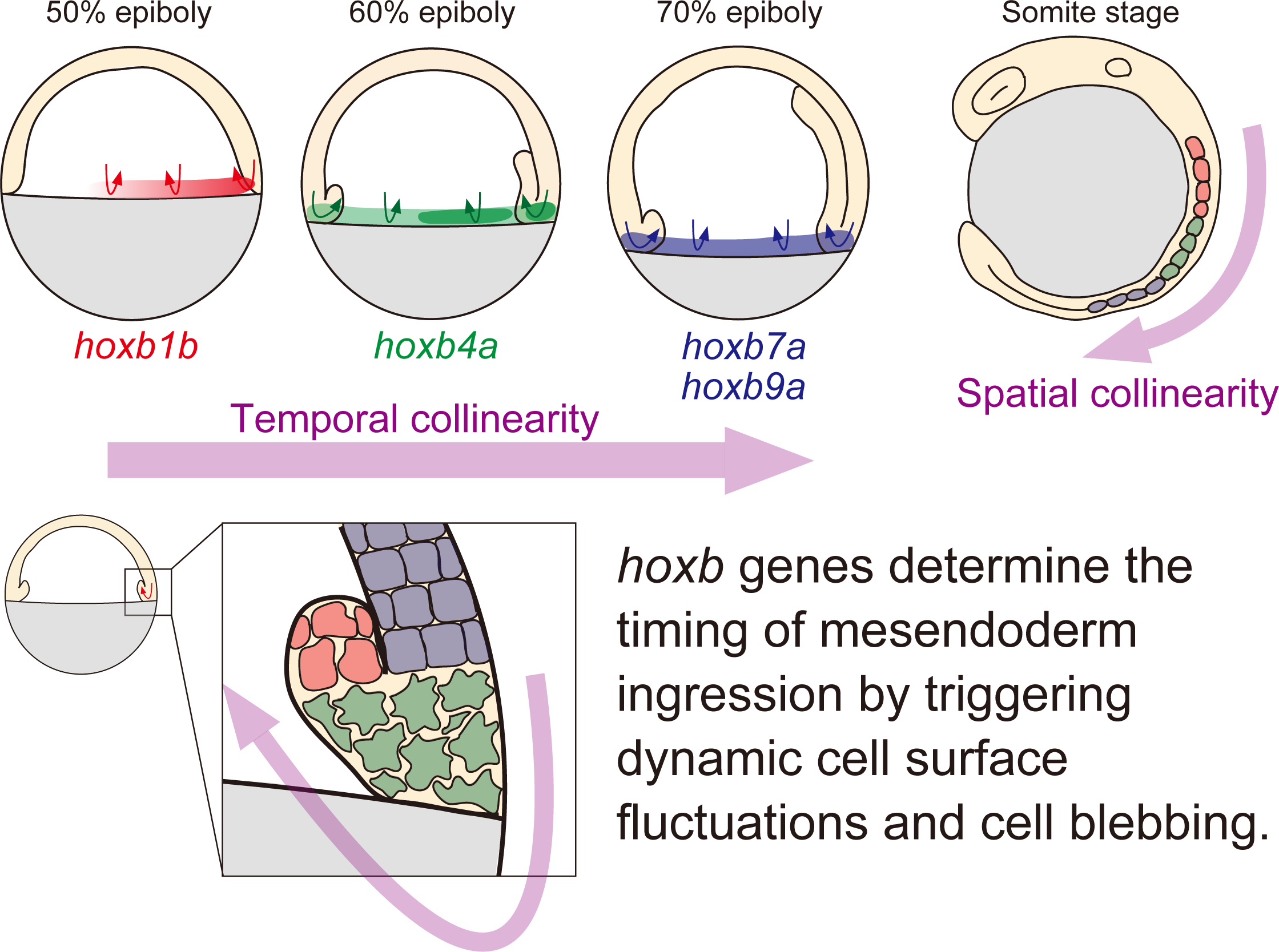

*hoxb* genes are expressed at the blastoderm margin in a temporally collinear manner and determine the timing of mesendoderm ingression by triggering dynamic cell surface fluctuations and cell blebbing. This results in a spatial collinearity of mesendoderm cell positioning along the anterior-posterior extent of the forming body axis with early ingressing cells being positioned more anteriorly than late ingressing cells.

## INTRODUCTION

The tight regulation of cell behaviors in time and space is essential for normal embryonic development. Gastrulation is the first morphogenetic process in embryonic development when dynamic and orchestrated cell behaviors lead to the formation of the three germ layers - ectoderm, mesoderm and endoderm. In zebrafish, gastrulation movements can be subdivided into three major processes ^1–4^: first, blastoderm cells spread over the yolk cell in a movement named ‘epiboly’. Second, mesendoderm progenitor cells at blastoderm margin internalize by synchronized cell ingression to form the mesendoderm cell layer (hypoblast) below the non-internalizing ectoderm progenitors (epiblast) ^5,6^. Third, cells within both the hypoblast and epiblast undergo convergence and extension movements leading to the formation and anterior-posterior elongation of the embryonic body axis. Synchronized mesendoderm cell internalization in zebrafish involves both cell autonomous and cell non-autonomous effects ^6^. We recently showed that mesendoderm progenitors can be subdivided into leader and follower cells with leader cells showing higher protrusive activity and cell-autonomously ingressing at the germ ring margin, while follower cells display lower protrusive activity and need to be pulled inside during ingression by the leader cells ^7^.

*hox* genes are transcription factors playing a pivotal role in determining anterior-posterior patterning in bilaterians ^8^. They are arranged in chromosomal gene clusters and their positional order from 3’ to 5’ translates into their expression domains along anterior-posterior body axis and the onset of their initial expression within the embryo, a phenomenon called “spatial collinearity” ^9,10^ and “temporal collinearity” ^11–14^ , respectively. Work on *hoxb* genes in gastrulating chick embryos has shown that these two collinearities are tightly linked: *hoxb* genes are expressed in a temporally ordered manner (temporal collinearity) and their expression in the epiblast lateral to the primitive streak regulates the timing of mesodermal cell ingression into the primitive streak, which again translates into their localization along the forming anterior-posterior body axis after the completion of gastrulation ^15^. This suggests that the spatial collinearity of *hoxb* genes is directly established by the temporal collinearity of their expression. Similarly, posterior *hox* genes (paralogues 9-13) in tail-bud stage chick embryos are expressed in a temporally collinear manner within the presomitic mesoderm, progressively slowing down body axis elongation by repressing Wnt signaling ^16^. In *Xenopus* embryos, *hox* genes are also expressed in temporally collinear manner during gastrulation, starting within the mesoderm and then expanding into the presumptive ectoderm. This led to the “time space translator model”, where the temporal collinear *hox* expression within the mesoderm and its interaction with the embryonic organizer (Spemann organizer) defines positional information along the anterior-posterior axis in mesoderm and overlying neuroectoderm ^17–21^. While these different studies suggest that *hox* genes regulate the timing of cell involution/ingression/migration during gastrulation, which again results in the positioning of cells along the anterior-posterior body axis, the mechanism(s) by which *hox* genes function in these processes is still insufficiently understood.

Cell blebbing is a characteristic cellular behavior, playing important roles in cell migration during development and disease. The formation of cellular blebs is triggered by the transient detachment of plasma membrane from the underlying actin cortex, leading to the formation of a spherical membrane protrusions through intracellular pressure ^22,23^. In zebrafish, primordial germ cells use cell blebbing for their directional migration in response to chemotaxis utilizing SDF-1a/CXCR4 ligand-receptor pairs ^24^. Similar cell behaviors were observed in primordial germ cells in *Drosophila* embryos ^25^, and tumour cells that utilize cell blebbing for their directional migration. Recently, cell blebbing and associated cell surface fluctuations have been implicated in cell sorting in early mouse embryos ^26^. However, the role of cell blebbing and cell surface fluctuations for cell movements during gastrulation is still not fully understood.

Here we have examined the expression and function of *hoxb* genes in zebrafish gastrulation. We found that *hoxb* genes are expressed in a temporally collinear manner at the blastoderm margin during gastrulation, and that this collinear expression determines the timing of mesendoderm progenitor cell ingression and thus their localization along the anterior-posterior extent of the forming body axis. *hoxb* genes appear to function in mesendoderm progenitor internalization by controlling their cell surface fluctuation and blebbing activity.

## RESULTS

### *hoxb* gene expression at the blastoderm margin exhibits temporal collinearity during gastrulation

To understand how *hox* genes function in zebrafish gastrulation, we first examined the expression patterns of *hoxb* genes during gastrulation. We focussed on *hoxb* genes as they had previously been shown to regulate the timing of cell ingression during chick gastrulation ^15^. Amongst the *hoxb* genes, we analyzed *hoxb1a*, *hoxb1b*, *hoxb4a*, *hoxb7a* and *hoxb9a* as representatives for anterior (*hoxb1a* and *hoxb1b*), middle (*hoxb4a*) and posterior (*hoxb7a* and *hoxb9a*) *hoxb* genes, similar to what has been used previously ^15^. While *hoxb1* has two paralogues, *hoxb1a* and *hoxb1b*, which might be duplicated by a teleost-specific whole genome duplication, *hoxb4*, *hoxb7* and *hoxb9* have only single paralogues. We found that while *hoxb1a* expression was largely absent during gastrulation until 90% epiboly stage (Figure 1Aa-Af, Figure S1E), *hoxb1b* expression was initiated at the dorsal blastoderm margin, except the dorsal most region, already at 50% epiboly stage (Figure 1Ba, arrowhead, 1F, red lines, Figure S1A) ^27^. This expression increased and expanded along the animal-vegetal and dorsal-ventral axes in epiblast cells around the blastoderm margin at shield (Figure 1Bb), eventually being expressed around the entire dorsal-ventral extent in both epiblast and hypoblast at 60 - 70% epiboly (Figure 1Bc, Bd). *hoxb1b* continued to be expressed within both epiblast and hypoblast until the end of gastrulation with slightly lower expression within its ventral portion (Figure 1Be, Bf, Figure S1F). *hoxb4a* started to be expressed within the blastoderm margin only at 60% epiboly stage with the highest expression levels in the dorsal most and dorsal-lateral regions of the margin (Figure 1Ca-Cc, arrowhead, 1G, red lines, Figure S1B). During subsequent stages of gastrulation, *hoxb4a* expression expanded along the animal-vegetal axis in both epiblast and hypoblast cells (Figure 1Cd-Cf, Figure S1G), with the anterior (animal) limit of its expression domain being positioned more posterior (vegetal) than that of *hoxb1b* (Figure 1Cf, Figure S1G). Finally, *hoxb7a* and *hoxb9a* only started to be expressed at the blastoderm margin at 70% epiboly (Figure 1Da-Dd, Ea-Ed, arrowheads, Figure S1C, D) and their expression lasted in epiblast and hypoblast cells until the end of gastrulation (Figure 1De, Df, Ee, Ef, Figure S1H, I). Notably, the anterior limit of their expression domains was positioned more posterior than that of *hoxb4a* (Figure 1Df, Ef). These spatiotemporal differences in the expression of *hoxb* genes gave rise to their spatial collinear expression pattern detectable within the epiblast and hypoblast at 90% epiboly, and neural tube and somites at the 6-somites stage (Figure 1Bf-Ef, Figure S1J-N). Taken together, the expression patterns of *hoxb* genes show temporal collinearity within the blastoderm margin during the course of gastrulation, leading to their spatial collinear expression within the trunk from late gastrulation onwards.

**Figure 1.**
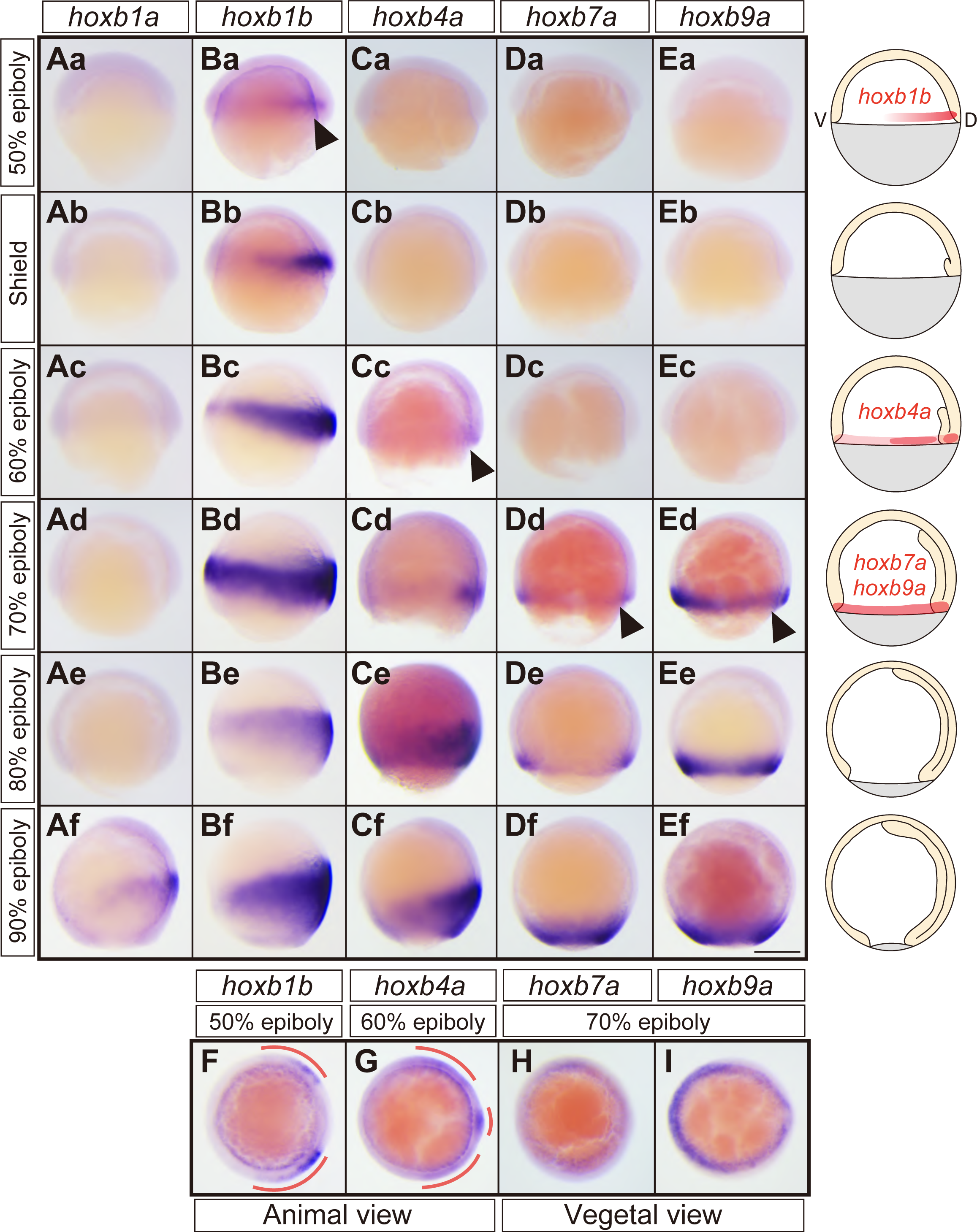
*hoxb* gene expression at blastoderm margin exhibits temporal collinearity during gastrulation. (A) Bright-field images (lateral views) of *hoxb1a* expression patterns at 50% epiboly (Aa), shield (Ab), 60% epiboly (Ac), 70% epiboly (Ad), 80% epiboly (Ae) and 90% epiboly (Af) stages. (B) Bright-field images (lateral views) of *hoxb1b* expression patterns at 50% epiboly (Ba), shield (Bb), 60% epiboly (Bc), 70% epiboly (Bd), 80% epiboly (Be) and 90% epiboly (Bf) stages. (C) Bright-field images (lateral views) of *hoxb4a* expression pattern at 50% epiboly (Ca), shield (Cb), 60% epiboly (Cc), 70% epiboly (Cd), 80% epiboly (Ce) and 90% epiboly (Cf) stages. (D) Bright-field images (lateral views) of *hoxb7a* expression pattern at 50% epiboly (Da), shield (Db), 60% epiboly (Dc), 70% epiboly (Dd), 80% epiboly (De) and 90% epiboly (Df) stages. (E) Bright-field images (lateral views) of *hoxb9a* expression pattern at 50% epiboly (Ea), shield (Eb), 60% epiboly (Ec), 70% epiboly (Ed), 80% epiboly (Ee) and 90% epiboly (Ef) stages. (F) Bright-field image (animal view) of *hoxb1b* expression pattern at 50% epiboly. (G) Bright-field image (animal view) of *hoxb4a* expression pattern at 60% epiboly. (H) Bright-field image (vegetal view) of *hoxb7a* expression pattern at 70% epiboly. (I) Bright-field image (vegetal view) of *hoxb9a* expression pattern at 70% epiboly. The rightmost column shows schematic diagrams for each of the developmental stages with the red color outlining the initial expression domains of *hoxb1b*, *4a*, *7a* and *9a* at the blastoderm margin. Arrowheads, initial expression of *hoxb* genes at the blastoderm margin during gastrulation. Dorsal side is to the right. Scale bar, 200 µm

### Loss-of-function of ‘early’ *hoxb* genes interferes with mesendoderm ingression and epiboly movements

Next we examined the function of *hoxb* genes during gastrulation. To this end, we first used *morpholino* oligonucleotides (MOs) blocking translation of *hoxb* genes (knock down, see STAR Methods for target sequences). *hoxb1a* MO injected embryos (*hoxb1a* morphants) showed no obvious defect during gastrulation compared to control MO-injected embryos, as expected from the absence of *hoxb1a* expression during gastrulation (Figure 2Aa-Ad, Ba-Bd). *hoxb1b* morphants, in contrast, exhibited diminished mesendoderm ingression and animal pole-directed migration at 6 hours post fertilization (hpf, shield stage) (Figure 2Ca, 2Da, red arrowhead), as evidenced by migrating mesendoderm cells being positioned closer to the germ ring and further away from the animal pole in *hoxb1b* morphants compared to control embryos at 7 hpf (60% epiboly, Figure 2Cb, 2Db, arrowheads). Notably, this reduced animal pole translocation of mesendoderm cells in 6 and 7 hpf *hoxb1b* morphants was detectable all around the germ ring including the most dorsal region where *hoxb1b* was not expressed (Figure 2Ca, red arrowhead, 2Cb, arrowhead). To determine whether this diminished translocation of mesendoderm cells applies to both mesoderm and endoderm cells, we analyzed endodermal cell positioning in *hoxb1b* morphants by visualizing the expression of the endodermal marker gene *sox17*. This showed that endoderm cell translocation towards the animal pole was reduced all around the germ ring in *hoxb1b* morphant embryos (Figure S2A, B, red arrowheads), suggesting that *hoxb1b* function is required for both mesoderm and endoderm cell ingression and animal pole-directed migration. Defective mesendoderm translocation in *hoxb1b* morphants at the onset of gastrulation was followed by reduced epiboly movements of the blastoderm towards the vegetal pole at 9 hpf (90% epiboly in control embryos) and 10.5 hpf (bud stage in control embryos) (Figure 2Cc, 2Cd, 2Dc and 2Dd, arrows).

In contrast to *hoxb1b* morphants, *hoxb4a*, *hoxb7a* and *hoxb9a* morphants did not exhibit any obvious mesendoderm ingression or animal pole-directed migration phenotype at 6 and 7 hpf (Figure 2 Ea-Ga, 2Eb-Gb), suggesting that they might be – at least partially – dispensable for these processes. Similar to *hoxb1b* morphants, however, *hoxb4a*, *hoxb7a* and *hoxb9a* morphants showed delayed blastoderm epiboly movements at the end of gastrulation (Figure 2Ec-Gc, 2Ed-Gd, arrows), suggesting a common function of *hoxb* genes in regulating blastoderm epiboly movements.

**Figure 2.**
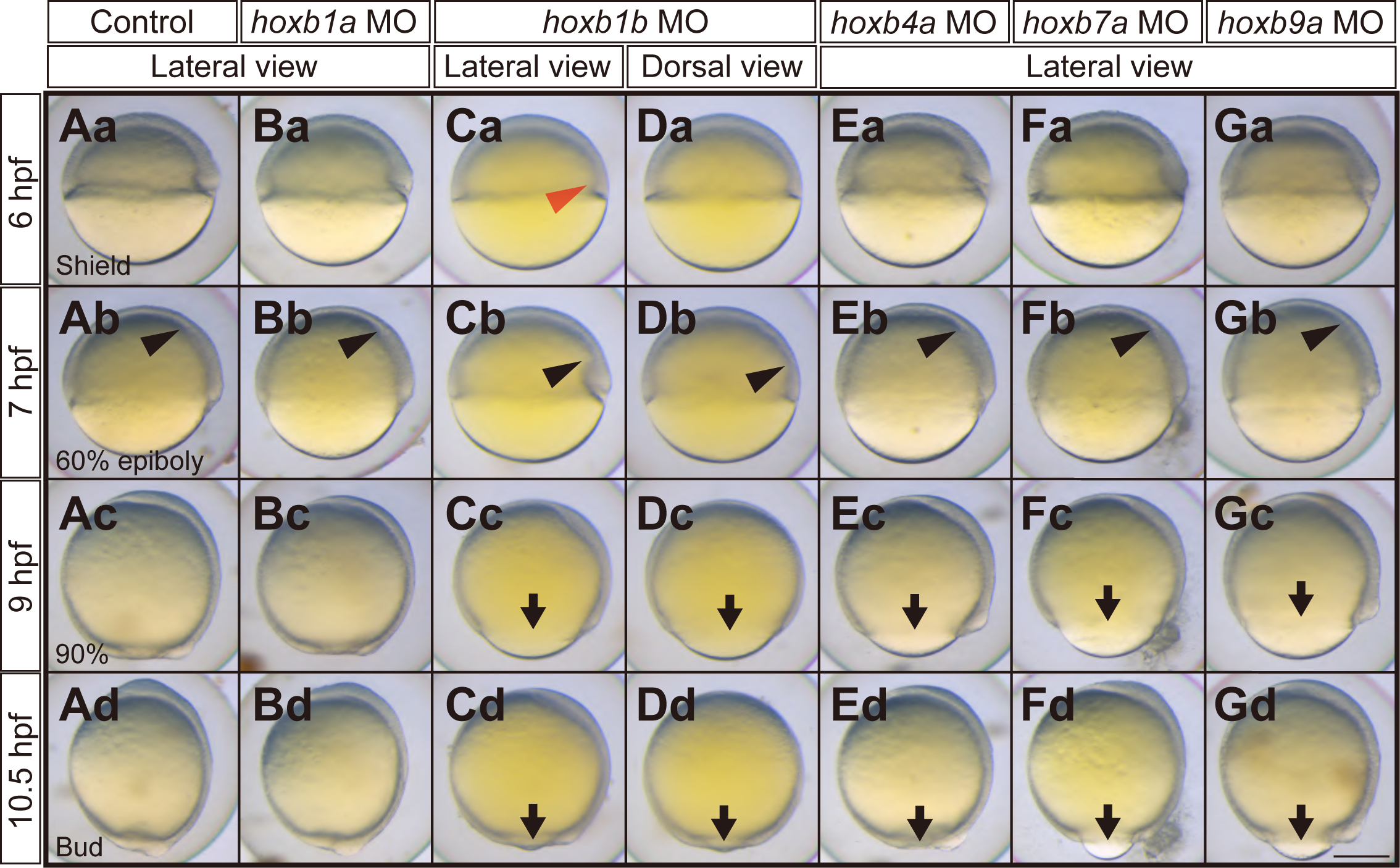
*hoxb* morphant embryos exhibit defective mesendodermal cell ingression/migration and epiboly movement delay. (A) Bright-field images (lateral view) of control antisense *morpholino* oligonucleotide (MO)-injected embryos at 6 hpf (shield, Aa), 7 hpf (60% epiboly, Ab), 9 hpf (90% epiboly, Ac), 10.5 hpf (bud, Ad). (B) Bright-field images (lateral view) of *hoxb1a* MO injected embryos (morphants) at 6 hpf (Ba), 7 hpf (Bb), 9 hpf (Bc), 10.5 hpf (Bd). (C, D) Bright-field images (lateral view, C and dorsal view, D) of *hoxb1b* morphants at 6 hpf (Ca, Da), 7 hpf (Cb, Db), 9 hpf (Cc, Dc), 10.5 hpf (Cd, Dd). (E) Bright-field images (lateral view) of *hoxb4a* morphants at 6 hpf (Ea), 7 hpf (Eb), 9 hpf (Ec), 10.5 hpf (Ed). (F) Bright-field images (lateral view) of *hoxb7a* morphants at 6 hpf (Fa), 7 hpf (Fb), 9 hpf (Fc), 10.5 hpf (Fd). (G) Bright-field images (lateral view) of *hoxb9a* morphants at 6 hpf (Ga), 7 hpf (Gb), 9 hpf (Gc), 10.5 hpf (Gd). Red arrowheads point at defective cell mesendoderm ingression at the dorsal blastoderm margin at 6 hpf. Black arrowheads point at the leading edge of mesendodermal cells migrating towards the animal pole. Arrows point at the blastoderm margin for the embryos exhibiting epiboly delay. Dorsal side is to the right. Scale bar, 200 µm.

To test the specificity of the observed *hoxb* morphant phenotypes, we also generated loss-of-function mutants for *hoxb1b, hoxb4a, hoxb7a* and *hoxb9a* genes by CRISPR/Cas9 (Figure S2D, see STAR Methods for target sequences). We found that *hoxb1b, hoxb4a, hoxb7a* and *hoxb9a* mutants embryos displayed phenotypes similar to their morphant counterparts (Figure S2E-J), supporting the specificity of the observed morphant phenotypes.

### Premature expression of ‘middle’ or ‘late’ *hoxb* genes interferes with mesendoderm ingression and epiboly movements

To further determine the activity of *hoxb* genes during gastrulation, we injected full-length mRNAs of each *hoxb* gene and examined their overexpression phenotypes. We found that *hoxb1a* and *hoxb1b* mRNA injected embryos exhibited no obvious phenotype during gastrulation compared to control embryos (Figure 3Aa-Ad, Ba-Bd, Ca-Cd). *hoxb4a* mRNA injected embryos, in contrast, showed diminished mesendoderm ingression and animal pole-directed migration at 6 and 7 hpf (Figure 3Da, Db, red and black arrowheads), followed by delayed epiboly movements at 9 and 10.5 hpf (Figure 3Dc, Dd, arrows). Similar phenotypes were also observed in embryos overexpressing *hoxb7a* or *hoxb9a* (Figure 3E, F), suggesting that *hoxb4a, hoxb7a* or *hoxb9a,* upon premature and ectopic expression, exhibit comparable activity in suppressing mesendoderm ingression and animal pole-directed migration during early gastrulation.

**Figure 3.**
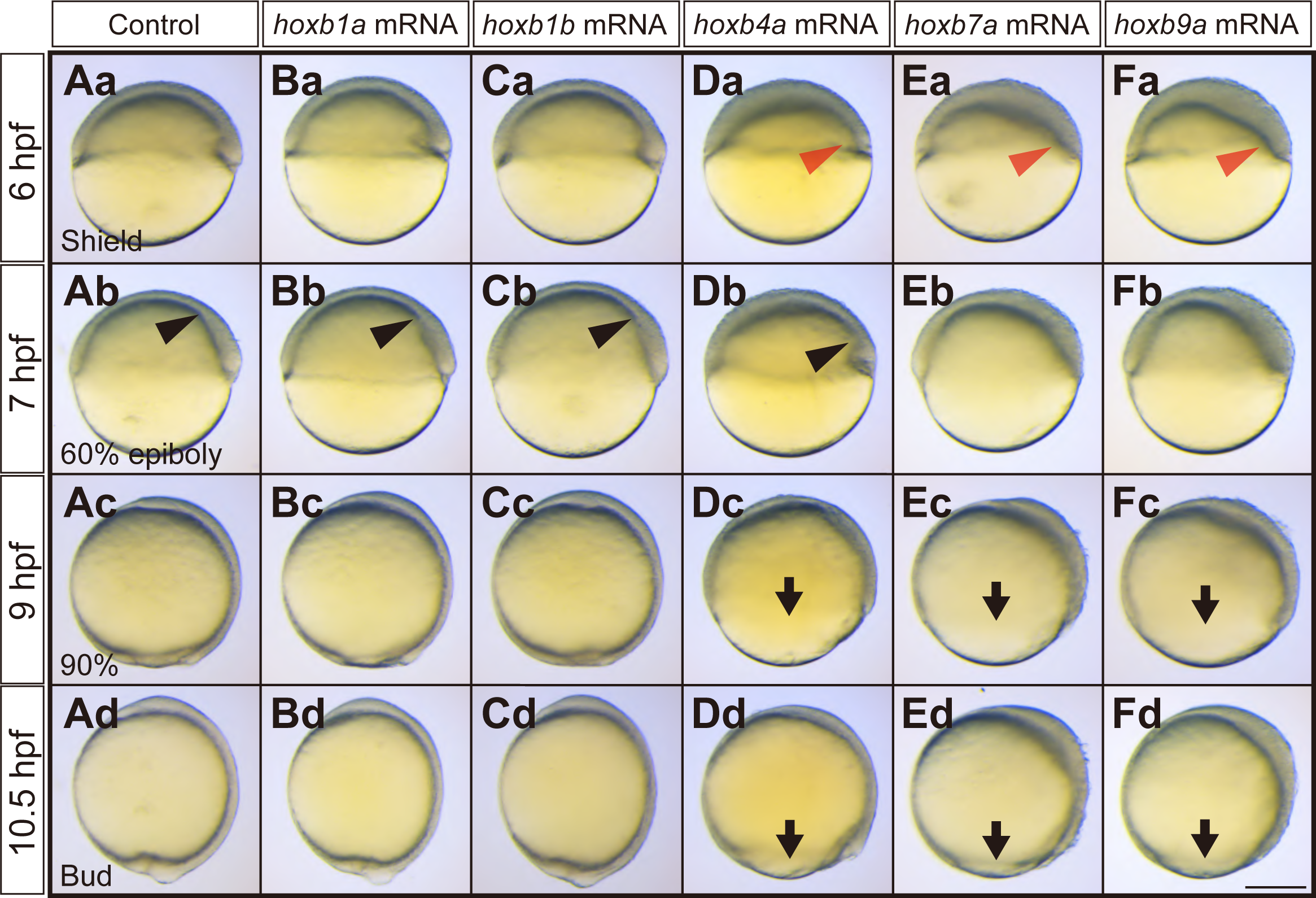
Middle or late *hoxb* overexpressed embryos exhibit defective mesendodermal cell ingression/migration and epiboly movement delay. (A) Bright-field images (lateral views) of water-injected control embryos at 6 hpf (Aa), 7 hpf (Ab), 9 hpf (Ac), 10.5 hpf (Ad). (B) Bright-field images (lateral views) of *hoxb1a* mRNA injected embryos at 6 hpf (Ba), 7 hpf (Bb), 9 hpf (Bc), 10.5 hpf (Bd). (C) Bright-field images (lateral views) of *hoxb1b* mRNA injected embryos at 6 hpf (Ca), 7 hpf (Cb), 9 hpf (Cc), 10.5 hpf (Cd). (D) Bright-field images (lateral views) of *hoxb4a* mRNA injected embryos at 6 hpf (Da), 7 hpf (Db), 9 hpf (Dc), 10.5 hpf (Dd). (E) Bright-field images (lateral views) of *hoxb7a* mRNA injected embryos at 6 hpf (Ea), 7 hpf (Eb), 9 hpf (Ec), 10.5 hpf (Ed). (F) Bright-field images (lateral views) of *hoxb9a* mRNA injected embryos at 6 hpf (Fa), 7 hpf (Fb), 9 hpf (Fc), 10.5 hpf (Fd). Red arrowheads point at defective mesendoderm ingression at dorsal blastoderm margin at 6 hpf. Black arrowheads point at the leading edge of mesendodermal cells migrating towards the animal pole. Arrows point at the blastoderm edge for the embryos exhibiting epiboly delay. Dorsal side is to the right. Scale bar, 200 µm.

To determine whether the observed mesendoderm phenotypes in *hoxb4a*, *hoxb7a* or *hoxb9a* overexpressing embryos were due to aberrant mesendoderm morphogenesis and not reduced mesendoderm specification, we analyzed the expression of the pan-mesendodermal marker gene *no tail* (*ntl*) in *hoxb4a*, *hoxb7a* or *hoxb9a* overexpressing embryos. We found that *ntl* expression levels at shield stage (6 hpf) did not show any major changes in *hoxb4a*, *hoxb7a* or *hoxb9a* mRNA injected compared to control embryos (Figure S3Aa-Da), suggesting that the observed mesendoderm defects in *hoxb4a*, *hoxb7a* or *hoxb9a* overexpressing embryos are due to aberrant morphogenesis and not changes in cell fate specification. According to the concept of ‘posterior prevalence’, postulating the hierarchical and functional dominance of posterior *hox* genes over more anterior *hox* genes ^12,28^, the phenotypes we observed here can be interpreted as that premature overexpression of ‘late’ or ‘posterior’ *hoxb* genes interfere with the function of ‘early’ or ‘anterior’ *hoxb* genes, regulating early mesendoderm cell ingression.

### *hoxb* genes determine the timing of mesendodermal cell ingression

*Hox* genes have previously been shown to determine the timing of mesoderm cell ingression during chick gastrulation ^15^. To investigate whether *hoxb* genes might have a similar function in zebrafish gastrulation, we analyzed mesendoderm ingression in *hoxb* loss or gain-of-function embryos. To this end, we labelled cells within the lateral blastoderm margin with fluorescent tracer DiI at the stages when each *hoxb* gene is initially expressed and traced the labelled cell populations in control and *hoxb* loss-of-function embryos (Figure 4A, E, I). In nearly all control embryos at 50% epiboly stage, when endogenous *hoxb1b* initiates its expression (Figure 1Ba), labelled mesendoderm progenitors had completed their migration into the hypoblast 30 minutes (min) after their labelling. In contrast, in only 10% of the *hoxb1b* morphant embryos, the labelled mesendoderm cells had ingressed 30 min after their labelling, and even after 60 min, only 80% of the morphant embryos showed complete ingression of labelled cells (Figure 4A, Ba-Bc, Ca-Cc, D, black and red arrowheads). This suggests that *hoxb1b* is required for timely ingression of mesendoderm cells at shield stage.

**Figure 4.**
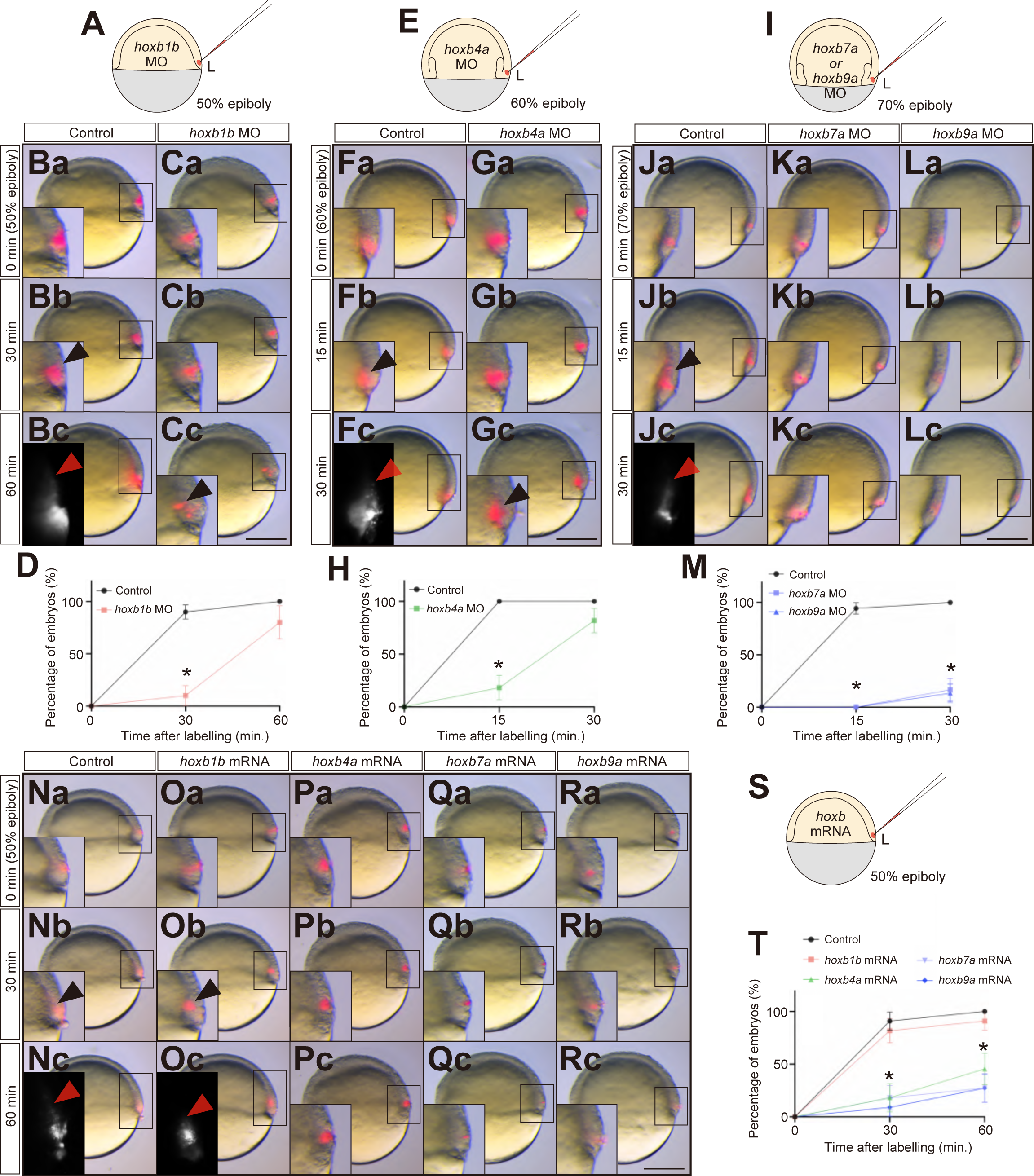
*hoxb* expression is required for proper mesendoderm morphogenesis. (A) Schematic representation of DiI cell labelling at the lateral blastoderm margin at 50% epiboly for *hoxb1b* morphants. (B, C) Bright-field/fluorescence images of labelled cells in control (B) and *hoxb1b* morphants (C) at 0 min (50% epiboly, just after labelling, a), 30 min (b) and 60 min after labelling (c). (D) Percentage of control (black) and *hoxb1b* morphant (red) embryos with all labelled cells having completed their ingression as a function of time after the labelling at 50% epiboly (n = 20 for control, n = 10 for *hoxb1b* morphants). (E) Schematic representation of DiI cell labelling at the lateral blastoderm margin at 60% epiboly for *hoxb4a* morphants. (F, G) Bright-field/fluorescence images of labelled cells in control (F) and *hoxb4a* morphants (G) at 0 min (60% epiboly, just after labelling, a), 15 min (b) and 30 min after labelling (c). (H) Percentage of control (black) and *hoxb4a* morphant (green) embryos with all labelled cells having completed their ingression as a function of time after the labelling at 60% epiboly (n = 12 for control, n = 11 for *hoxb4a* morphants). (I) Schematic representation of DiI cell labelling at the lateral blastoderm margin at 70% epiboly for *hoxb7a* or *hoxb9a* morphants. (J-L) Bright-field/fluorescence images of labelled cells in control (J), *hoxb7a* morphants (K) and *hoxb9a* morphants (L) at 0 min (70% epiboly, just after labelling, a), 15 min (b) and 30 min after labelling (c). (M) Percentage of control (black) and *hoxb7a* (grey*) and hoxb9a* (blue) morphant embryos with all labelled cells having completed their ingression as a function of time after the labelling at 70% epiboly (n = 18 for control, n = 12 for *hoxb7a* morphants, n = 15 for *hoxb9a* morphants). (N-R) Bright-field/fluorescence images of labelled cells in control (N), *hoxb1b* (O), *hoxb4a* (P), *hoxb7a* (Q) and *hoxb9a* (R) mRNA injected embryos at 0 min (50% epiboly, just after labelling, a), 30 min (b) and 60 min after labelling (c). (S) Schematic representation of DiI cell labelling at the lateral blastoderm margin at 50% epiboly for *hoxb* mRNA injected embryos. (T) Percentage of control (black), *hoxb1b* (red), *hoxb4a* (green), *hoxb7a* (grey) or *hoxb9a* (blue) mRNA injected embryos with all labelled cells having completed their ingression as a function of time after the labelling at 50% epiboly (n = 11 for control, n = 11 for *hoxb1b* mRNA, n = 11 for *hoxb4a* mRNA, n = 11 for *hoxb7a* mRNA, n = 11 for *hoxb9a* mRNA). *p<0.05. Values are reported as mean±s.e.m. and p values were determined by the *t*-test. Black arrowheads point at ingressed mesendodermal cells. Red arrowheads point at the leading edge of the ingressed mesendoderm. Ventral view with lateral (left side) to the right. Insets are magnified views of the boxed areas in the lower magnification images. Scale bar, 200 µm.

In *hoxb4a* morphants at 60% epiboly, corresponding to the stage when *hoxb4*a is initially expressed (Figure 1Cc), ingression of labelled cells was severely reduced with only in 18% of the labelled embryos showing complete cell ingression 15 min after the labelling, a stage when labelled cell ingression was completed in all of the control embryos (Figure 4E, Fa-Fc, Ga-Gc, H, black and red arrowheads). Finally, *hoxb7a* or *hoxb9a* morphants at 70% epiboly, when these genes begin to be expressed within the blastoderm margin (Figure 1Dd, Ed), also showed strongly reduced ingression of labelled cells, resulting in only 16% and 10% of the morphant embryos, respectively, showing complete cell ingression 30 min after labelling, when labelled cell ingression was completed in all control embryos (Figure 4I, Ja-Jc, Ka-Kc, La-Lc, M, black and red arrowheads). These results indicate that *hoxb* genes are required for punctual ingression of mesendoderm cells at the timepoint of their initial expression.

To test whether *hoxb* gene expression is also sufficient to control the timing of mesendoderm cell ingression, we analyzed cell ingression at shield stage (6 hpf) in embryos overexpressing different *hoxb* genes (Figure 4S). While overexpression of *hoxb1b* had no major effect on labelled mesendoderm cell ingression at shield stage (Figure 4Na-Nc, Oa-Oc, T), embryos overexpressing *hoxb* genes, the endogenous expression of which is only initiated at later stages of gastrulation (*hoxb4a*, *hoxb7a* and *hoxb9a*. Figure 1Cc, Dd, Ed), exhibited clearly reduced cell ingression at shield stage (Figure 4Pa-Pc, Qa-Qc, Ra-Rc, T). This suggests that ectopic and premature expression of ‘late’ *hoxb* genes (*hoxb4a, hoxb7a* and *hoxb9a*) is sufficient to interfere with mesendoderm cell ingression at the onset of gastrulation, and this is also consistent with the concept of ‘posterior prevalence’.

To analyze which aspects of mesendoderm cell ingression are altered by *hoxb* over- and under-expression, we traced cell ingression in control, *hoxb1b* morphant and *hoxb7a* overexpressing embryos at the onset of gastrulation (6 hpf, Figure 2Ca, Figure 3Ea). For visualizing and tracing cells in 3 dimensions over time (4D), we injected dextran mini-ruby into the intercellular space and recorded mesendoderm cell ingression within the lateral germ ring (Figure 5A, described as ‘0 min’ in Figure 5B, I, P). In control embryos, cells first moved vegetally towards the germ ring margin, followed by ingression towards the yolk cell and migration away from the germ ring margin towards the animal pole of the gastrula (Figure 5B-F). Consistent with our previous observations ^7^, we found that cells right at the germ ring margin were the first undergoing ingression and migration away from the margin towards the animal pole, sequentially followed by cells positioned further away from the margin first moving towards the margin and then undergoing ingression (Figure 5B-H, yellow, red, green arrowheads ^7^). In *hoxb1b* morphant and *hoxb7a* overexpressing embryos, mesendoderm cell ingression occurred in a similar pattern as in control embryos. However, the onset of cell ingression was delayed (Figure 5I-W, yellow, red, green arrowheads) and the velocity of cell movement during ingression was significantly reduced (Figure 5X), suggesting that spatiotemporally ordered expression of *hoxb* genes is required for timely and efficient mesendoderm cell ingression at the onset of gastrulation.

**Figure 5.**
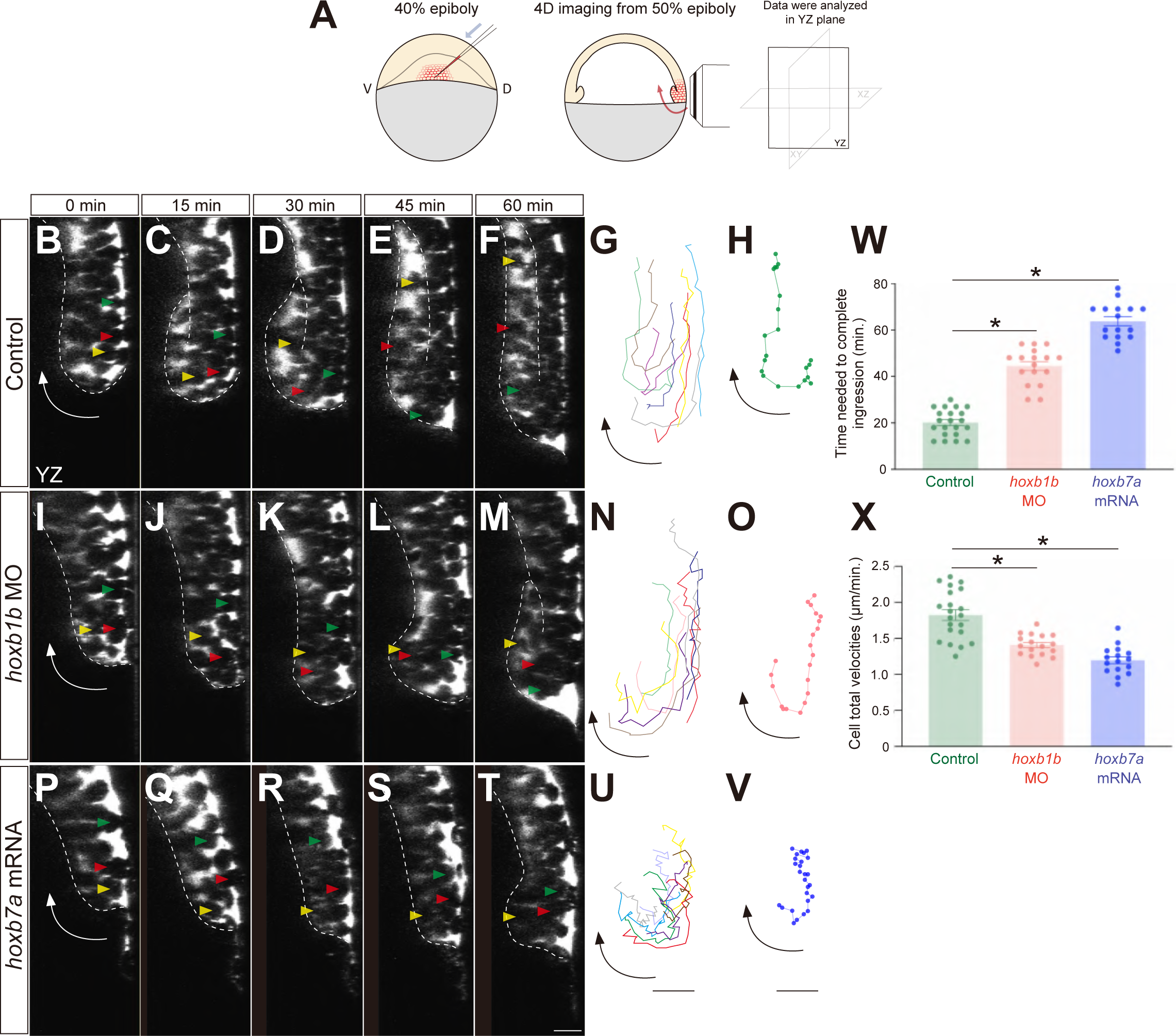
*hoxb* expression determines the timing of mesendodermal cell ingression. (A) Schematic representation of mesendodermal cell movement analysis. Embryos were injected with dextran mini-ruby to label the intercellular space (interstitial fluid) and cells at the blastoderm margin were identified by the absence of labelling and tracked in 4D at the lateral side of the embryo from 50% epiboly stage onwards. D, dorsal. V, ventral. (B-F, I-M, P-T) Confocal fluorescence images of the lateral blastoderm margin at consecutive stages of mesendodermal cell ingression from 50% epiboly stage (0 min) onwards in control embryos (B-F), *hoxb1b* morphant (I-M) and *hoxb7a* overexpressing embryos (P-T). Arrows depict the overall movement direction of cells at the blastoderm margin. Yellow, red and green arrowheads depict individual mesendodermal cells during ingression. Scale bar, 20 µm. (G, N, U) Representative tracks of mesendodermal cells undergoing ingression in control (G), *hoxb1b* morphant (N) and *hoxb7a* overexpressed embryos (U). 8 cells were tracked in each condition. Cells were tracked for 0-60 minutes (3 minutes/frame) in control embryos and *hoxb1b* morphants, and for 0-75 minutes (3 minutes/frame) in *hoxb7a* overexpressing embryos. Scale bar, 20 µm. (H, O, V) Representative single cell track of a mesendodermal cell undergoing ingression at the blastoderm margin at 50% epiboly in control (H), *hoxb1b* morphant (O) and *hoxb7a* overexpressed embryos (V). Dots represents each time points. Scale bar, 20 µm. (W) Average time needed for mesendodermal cells to complete ingression from 50% epiboly onwards in control (n = 21, N = 3), *hoxb1b* morphants (n = 17, N = 3) and *hoxb7a* overexpressing embryos (n = 16, N = 3). Mean±s.e.m; *p<0.05 (*t*-test). (X) Average cell total velocities of mesendodermal cells undergoing ingression in control (n = 21, N = 3), *hoxb1b* morphant (n = 17, N = 3) and *hoxb7a* overexpressing embryos (n = 16, N = 3). Mean±s.e.m; *p<0.05 (*t*-test).

### *hoxb* genes regulate mesendodermal cell ingression timing at the blastoderm margin in a cell-autonomous manner

To examine how *hoxb* genes function in mesendoderm cell ingression, we transplanted ∼ 10-20 control and *hoxb1b* or *hoxb7a* morphant or overexpressing cells from the lateral germ ring margin of donor embryos into the corresponding region of a control host embryo at 40% epiboly stage. We then traced the ingression behavior of the transplanted cells in 4D and determined the timing of cell ingression for each of the transplanted cells (Figure 6A). When only control cells were transplanted, the transplanted cells ingressed in close succession with an average time-difference of ∼ 2 minutes (Figure 6B, E, Video S1). In contrast, when control cells were co-transplanted with *hoxb1b* morphant or *hoxb7a* overexpressing cells, ingression of the morphant/ overexpressing cells was delayed by ∼ 52 min compared to control cells (Figure 6C, D, E, Video 2 and 3). Such delay was not observed when control cells were co-transplanted with *hoxb1b* overexpressing or *hoxb7a* morphant cells (Figure 6E, Figure S4A, B). This suggests that interfering with the expression of ‘early’ *hoxb* genes (*hoxb1b*) or premature expression of ‘late’ *hoxb* genes (*hoxb7a*) leads to a cell-autonomous delay in cell ingression at shield stage.

**Figure 6.**
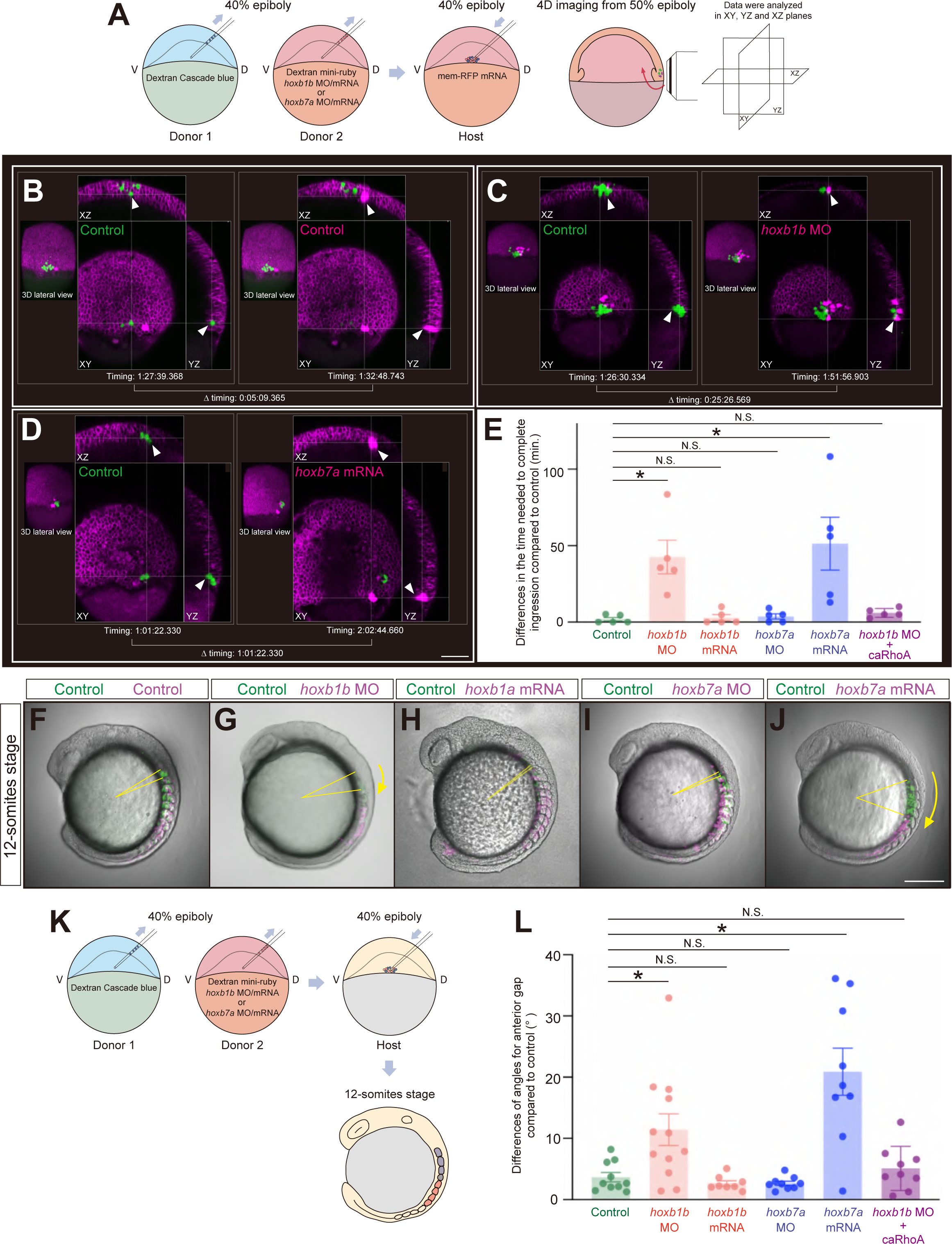
*hoxb* expression cell-autonomously determines the timing of mesendodermal cell ingression and their resulting position along the anterior-posterior body axis. (A) Schematic representation of the cell transplantation assay and subsequent 4D imaging. Cells aspirated from the lateral blastoderm margin of embryos injected with dextran cascade blue (donor 1) and a combination of dextran mini-ruby and *hoxb1b* MO/mRNA or *hoxb7a* MO/mRNA (donor 2) were transplanted into the corresponding region of a host embryo injected with membrane-RFP mRNA. The movement of the transplanted cells was monitored in 4D from 50% epiboly stage onwards. D, dorsal. V, ventral. (B-D) Time needed for the transplanted cells to complete ingression in host embryos containing a combination of transplanted control cells (green, left) with control (B), *hoxb1b* MO (C) or *hoxb7a* mRNA(D) cells (magenta, right). Scale bar, 100 µm. (E) Average difference in the time needed for transplanted control cells and co-transplanted control (n = 5), *hoxb1b* MO (n = 5), *hoxb1b* mRNA (n = 5), *hoxb7a* MO (n = 5), *hoxb7a* mRNA (n = 5), or *hoxb1b* MO plus caRhoA mRNA (n = 5) injected cells to complete ingression. Arrowheads point at ingressing mesendodermal cells. Mean±s.e.m; *p<0.05 (*t*-test); N.S., not significant. (F-J) Localization of transplanted cells along the anterior-posterior body axis within the somitic mesoderm of host embryos at the 12-somite stage for control (green) co-transplanted with control (F), *hoxb1b* MO (G), *hoxb1* mRNA (H), *hoxb7a* MO (I) or *hoxb7a* mRNA (J) injected cells (magenta) at 40% epiboly stage. Yellow arrows outline the angles between the most anteriorly located control cells (green) and co-transplanted control, *hoxb1b* MO/mRNA or *hoxb7a* MO/mRNA cells (magenta). Scale bar, 200 µm. (K) Schematic representation of double transplantation assay for determining distribution patterns of transplanted cells in 12-somite stage embryos. (L) Average angles between the most anteriorly located control cells and co-transplanted control (n = 10), *hoxb1b* MO (n = 12), *hoxb1b* mRNA (n = 8), *hoxb7a* MO (n = 9), *hoxb7a* mRNA (n = 9) or *hoxb1b* MO plus caRhoA mRNA (n = 9) injected cells within the somitic mesoderm of host embryos at the 12-somite stage. Mean±s.e.m; *p<0.05 (*t*-test); N.S., not significant.

### *hoxb*-mediated timing of mesendodermal cell ingression during gastrulation translates into their localization along anterior-posterior body axis post gastrulation

To determine how the timing of mesendoderm cell ingression translates into the positioning of the ingressed cells along the forming body axis, we performed the same transplantation assay as described above and analyzed the distribution pattern of the transplanted cells at 12-somites stage (after completion of gastrulation) (Figure 6K). We found that delayed ingression of *hoxb1b* morphant or *hoxb7a* overexpressing cells led to these cells being positioned more posteriorly along the body axis than the co-transplanted control cells (Figure 6F, G, J, L). No such more posterior positioning was found when control cells were co-transplanted with *hoxb1b* overexpressing or *hoxb7a* morphant cells (Figure 6H, I, L). Taken together, these results suggest that *hoxb* genes determine the proper timing for mesendoderm cell ingression, and that the timing of cell ingression determines the positioning of the ingressed cells along the anterior-posterior axis of the forming body axis.

### Initiation of frequent mesendodermal cell blebbing at blastoderm margin regulated by *hoxb* gene expression triggers cell ingression

To obtain some initial insight into the mechanisms by which *hoxb* genes determine the timing of mesendoderm cell ingression, we analyzed the protrusive behavior of ingressing cells, which we previously showed to mediate their cell ingression behavior ^7^. We reasoned that *hoxb* genes might control the timing of cell ingression by promoting the protrusive activity of mesendoderm cells. To challenge this possibility, we performed the same transplantation assay as described above and visualized the protrusive activity of the transplanted mesendoderm cells. We found that the number of F-actin-filled protrusions toward yolk syncytial layer formed by control and *hoxb1b* morphant or *hoxb7a* overexpressing cells was not significantly different (Figure S5A-E), suggesting that the formation of actin-rich cell protrusions as such is not regulated by *hoxb* genes.

Given that cell blebbing behavior has previously been observed within the blastoderm margin during zebrafish gastrulation ^29^ and recent studies have shown that cell blebbing and surface fluctuations play pivotal roles for segregation of epiblast and primitive endoderm in early mammalian embryos ^26^, we asked whether *hoxb* genes might determine the timing of mesendoderm cell ingression by upregulating blebbing/ surface fluctuations of mesendoderm cells. To address this possibility, we used the same transplantation assay as described above and examined the frequency of bleb formation in the transplanted mesendoderm cells (Figure 7A). When control cells were co-transplanted with *hoxb1b* morphant or *hoxb7a* overexpressing cells, control cells started to form frequent blebs shortly before initiating ingression; *hoxb1b* morphant or *hoxb7a* overexpressing cells, in contrast, started to form blebs later than the control cells (Figure 7B-E, arrowheads, Video 4-7). To determine the time-point when transplanted cells exhibited frequent blebs (defined as the state when cells form more than 5 blebs within 3 min), we monitored cell blebbing in control and *hoxb1b* morphant or *hoxb7a* overexpressing cells. We found that *hoxb1b* morphant and *hoxb7a* overexpressing cells exhibited frequent blebs significantly later than the control cells (Figure 7F). To test whether *hoxb*-dependent mesendoderm cell blebbing is an expression of increased cell surface fluctuations in mesendoderm cells, we quantified the dynamics of cell surface fluctuations in cells within the germ ring margin showing high versus low blebbing activity (Figure S5F-I). We found that cells exhibiting frequent blebbing directly preceding their ingression showed significantly larger cell surface fluctuations than non-ingressing cells exhibiting less blebbing (Figure 7G). This suggests that *hoxb* genes determine the timing of mesendoderm cell ingression by upregulating surface fluctuations in these cells.

**Figure 7.**
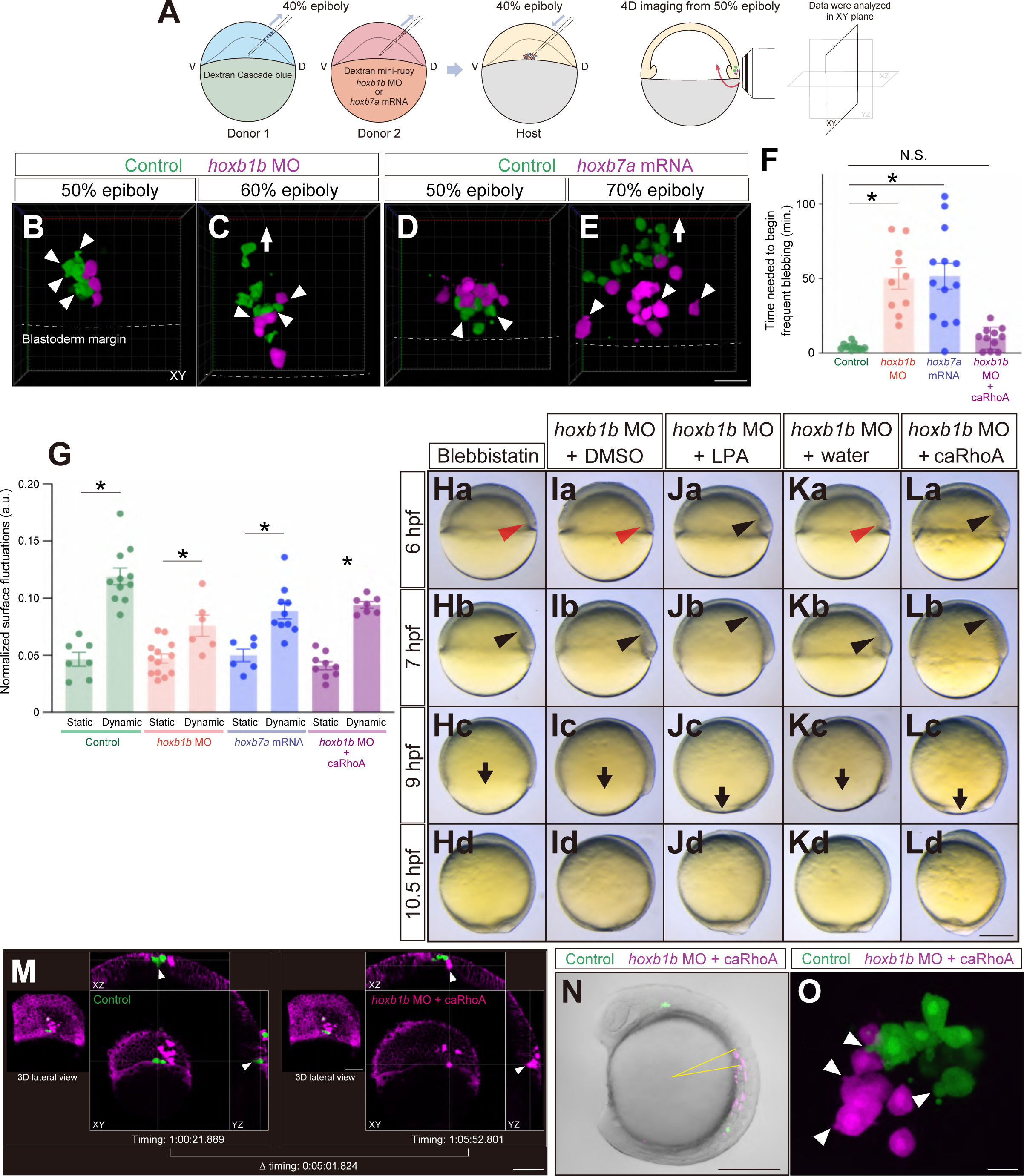
*hoxb* expression affects mesendodermal cell blebbing and associated cell surface fluctuations. (A) Schematic representation of the cell transplantation assay and subsequent 4D imaging. Cells aspirated from the lateral blastoderm margin of embryos injected with dextran cascade blue (donor 1) and a combination of dextran mini-ruby and *hoxb1b* MO or *hoxb7a* mRNA (donor 2) were transplanted into the corresponding region of a host embryo. The transplanted cell dynamics were monitored in 4D from 50% epiboly stage onwards. D, dorsal. V, ventral. (B-E) Confocal fluorescence images (3D projection) of transplanted control (green) and *hoxb1b* MO (B, C) or *hoxb7a* mRNA (D, E) injected cells (magenta) at 50% epiboly (B, D) and 60% (C) or 70% epiboly (E). Arrowheads depict cellular blebs. Arrow demarcates the migration direction of ingressed mesendodermal cells towards the animal pole. Dashed lines depict the borders between blastoderm margin and yolk. Scale bar, 60 µm (F) Time needed for mesendodermal cells to begin frequent blebbing after the onset of imaging in control (n = 12, N = 4), *hoxb1b* MO (n = 10, N = 3), *hoxb7a* mRNA (n = 13, N =4) or *hoxb1b* MO plus caRhoA mRNA (n = 11, N = 4) injected cells. Mean±s.e.m.; *p<0.05 (*t*-test). (G) Normalized surface fluctuations of control, *hoxb1b* MO, *hoxb7a* mRNA and *hoxb1b* MO plus caRhoA mRNA injected cells. Cells were analyzed before and after entering their frequent blebbing state (described as ’Static’ and ’Dynamic’, respectively). Mean±s.e.m.; *p<0.05 (*t*-test). (H-L) Phenotypes of Blebbistatin-treated embryos (H) or DMSO-treated (I), and LPA-treated (J), water-injected (K) or caRhoA mRNA injected (L) *hoxb1b* morphant embryos at 6 hfp (a), 7 hpf (b), 9 hpf (c) and 10.5 hpf (d). Red arrowheads point at defective mesendodermal cell ingression at the dorsal blastoderm margin at 6 hpf. Black arrowheads point at the leading edge of mesendodermal cells migrating towards the animal pole. Arrows point at the blastoderm margin at 9 hpf. Dorsal side is to the right. Scale bar, 200 µm. (M) Double transplantation assay for determining the ingression timing of control (green) and *hoxb1b* MO plus caRhoA mRNA injected (magenta) mesendodermal cells. Arrowheads indicate ingressing mesendodermal cells. For quantification and statistical analysis, see Figure 6E. Scale bars, 100 µm. (N) Distribution patterns of transplanted control (green) and *hoxb1b* MO plus caRhoA mRNA injected (magenta) mesendodermal cells in 12-somite stage embryo. Yellow line indicates the angles between the most anteriorly located control and *hoxb1b* MO plus caRhoA mRNA injected transplanted cells. For quantification and statistical analysis, see Figure 6L. Scale bar, 200 µm. (O) Blebbing behavior of transplanted control (green) and *hoxb1b* MO plus caRhoA mRNA injected (magenta) cells at 50% epiboly stage. For quantification and statistical analysis, see Figure 7F and 7G. Arrowheads depict cellular blebs. Scale bar, 20 µm.

Finally, to test whether increased cell blebbing and surface fluctuations constitutes the mechanism by which *hoxb* genes regulate the timing of mesendoderm cell ingression, we tested whether interfering with cell blebbing affects timely cell ingression. When we treated embryos with 10 μM Blebbistatin, an inhibitor of non-muscle myosin II reducing bleb formation, from 30 – 40% epiboly, we found that Blebbistatin-treated embryos, similar to *hoxb1b* morphant embryos, displayed delayed mesendoderm ingression and animal pole-directed migration at shield stage, and slower epiboly movements (Figure 7Ha-Hd, Figure 2Ca-Cd). To test whether increasing cell blebbing and surface fluctuations can also rescue the mesendoderm cell ingression phenotype in *hoxb1b* morphant embryos, we increased cell blebbing in *hoxb1b* morphants by treating them with lysophosphatidic acid (LPA) or overexpressing a constitutively active version of RhoA (caRhoA), which have previously been shown to increase actomyosin contractility and cell blebbing in zebrafish embryos ^30,31^. *hoxb1b* morphants treated with 10 μM LPA or injected with 1 pg caRhoA mRNA exhibited normal mesendoderm cell ingression and epiboly movements while DMSO-treated or water-injected embryos did not (Figure 7Ia-Id, Ja-Jd, Ka-Kd, La-Ld, Figure 2Aa-Ad). Notably, this rescue of cell movements in caRhoA mRNA injected *hoxb1b* morphant embryos phenotype was accompanied by normalized mesendoderm cell blebbing behavior, surface fluctuations, ingression timing, and distribution pattern along the anteroposterior axis at somite stage (Figure 6E, L, Figure 7F, G, M-O), further supporting the notion that *hoxb* genes control the timing of mesendoderm cell ingression by regulating cell blebbing and surface fluctuations.

## DISCUSSION

Our study demonstrates temporal collinearity of *hoxb* gene expression during zebrafish gastrulation. In line with this finding are previous reports on the temporal expression pattern of other *hox* genes, such as *hoxc4a* ^32^, *hoxb6b*, *hoxc6a* and *hoxc6b* ^32–34^, *hoxc8a* ^32,33,35^ and *hoxa9a*, *hoxb10a* and *hoxc10a* during zebrafish epiboly ^32,33^, showing that ‘anterior’ *hox* genes are expressed earlier than ‘posterior’ ones. Furthermore, *hox* genes in other species, such as mouse ^36,37^, chicken ^13,15,38–40^ , *Xenopus* ^17^ and cnidarians ^41^, have also been shown to exhibit temporal collinearity in their expression during gastrulation, demonstrating that temporal collinearity is a wide-spread feature of *hox* gene expression. Notably, however, in acoels ^42^ and annelids ^43^, and most recently also in *Xenopus tropicalis* ^44^, *hox* genes do not show temporally collinear expression, suggesting that temporal collinearity is not universally conserved in animal evolution.

Previous studies have shown that the temporal collinearity of *hoxb* gene expression regulates the timing of mesoderm cell ingression in the primitive streak during chick gastrulation ^15^. Moreover, this temporal collinearity of *hoxb* gene expression and mesoderm cell ingression in chicken translates into a spatial collinearity in mesoderm cell positioning along the anterior-posterior axis after gastrulation ^16^, important for limb positioning ^40^. Our present findings are consistent with those previous findings, indicating that *hoxb* gene function during gastrulation is conserved between teleosts and amniotes.

Importantly, our study goes beyond these previous observations by providing insight into the cell-autonomous and non-autonomous processes by which *hox* genes regulate mesendoderm cell ingression. In particular, our finding that under-expression of ‘early’ *hoxb* genes and over-expression of ‘late’ ones delay cell ingression in a cell-autonomous manner, suggests that *hox* genes determine cell ingression in a cell-autonomous manner ^45,46^. Yet, mesendodermal cell ingression has also been shown to display cell-non-autonomous features with ingression-competent cells taking along incompetent cells during ingression ^5^ and endoderm internalization being achieved by the collective internalization of mesodermal cells ^6^. Furthermore, our own recent findings suggest that ingressing mesendoderm cells can be subdivided into ‘leader’ cells, showing high protrusive activity and being able to cell-autonomously ingress at the onset of gastrulation, and ‘follower’ cells that are less protrusive and pulled inside by leader cells during ingression ^7^. Our present findings of early *hoxb* gene expression initiating the onset of mesendoderm ingression suggest that early mesendoderm ingression is a cell-autonomous process dependent on the timely expression of ‘early’ *hox* genes, such as *hoxb1b*.

Interestingly, our data also show that the premature expression of ‘late’ *hoxb* genes, such as *hoxb7a*, suppresses early mesendoderm cell ingression, suggesting that ‘late’ *hoxb* genes cannot trigger early mesendoderm ingression. How different *hox* genes control mesendoderm cell ingression in a stage-dependent manner is not yet clear, but it is conceivable that different *hox* genes elicit distinct cell behaviors required for cell ingression at different stages of gastrulation. Our finding that cells shortly before ingression display enhanced blebbing and surface fluctuations, point at the possibility that *hox* genes determine the timing of mesendodermal cell ingression by promoting cell surface fluctuations. This is consistent with previous studies showing that cell surface fluctuations play a pivotal role for segregation of epiblast and primitive endoderm in early mammalian embryos ^26^. Given that cell surface fluctuations have been linked to tissue fluidity ^26^, and that actively ingressing mesendoderm progenitor cells have been proposed to undergo motility-driven unjamming ^7^, it is conceivable that *hoxb* genes drive mesendoderm cell ingression by triggering their motility-driven unjamming. Yet, how the induction of such general behavior is modulated so that different *hox* genes can exert their function in a stage-specific manner remains to be investigated.

Our study suggests that *hoxb* genes determine mesendoderm cell ingression timing by upregulating RhoA/Rock-mediated cortical contractility and cell blebbing. However, this appears different from chicken gastrulation where mesendoderm cell ingression is initiated by basement membrane (BM) breakdown, which again is controlled by a loss of basally localized RhoA activity ^47^. Furthermore, blocking RhoA activity can rescue the delay in mesendoderm cell ingression in chicken embryos overexpressing *Hoxa13* ^16^, suggesting that *hox* genes determine the timing of mesendoderm cell ingression in chicken gastrulation by downregulating RhoA activity, rather than upregulating it as suggested by our present findings in zebrafish. How the molecular and cellular mechanisms by which *hox* genes determine the timing of mesendoderm ingression have been adapted to the specific organismal context during evolution remains an intriguing question for future research.

**Figure S1.**
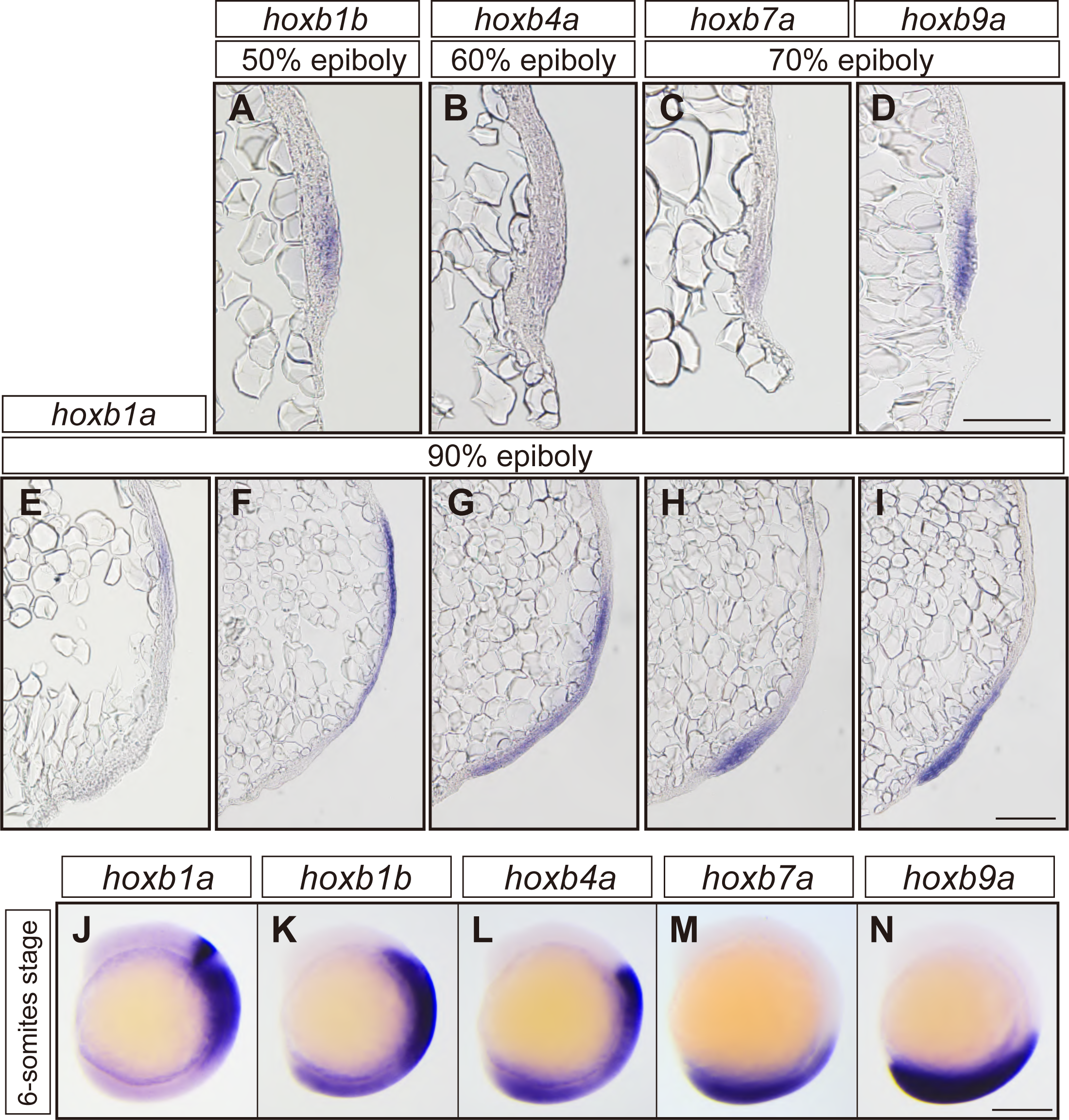
*hoxb* gene expression during gastrulation and somite stage - Related to Figure 1. (A-D) Bright-field images of cryosections of whole mount *in situ* hybridization in embryos showing *hoxb1b* at 50% (A), *hoxb4a* at 60% epiboly (B), *hoxb7a* at 70% epiboly (C), and *hoxb9a* at 70% epiboly (D). Only A is a tilted lateral view for capturing *hoxb1b* expression, and the others are lateral views. Scale bar, 20 µm. (E-I) Bright-field images (lateral views) of cryosections of whole mount *in situ* hybridization in embryos showing *hoxb1a* (E)*, hoxb1b* (F)*, hoxb4a* (G), *hoxb7a* (H) *and hoxb9a* (I) at 90% epiboly. Scale bar, 20 µm. (J-N) Bright-field images (lateral views) of expression patterns of *hoxb1a* (J), *hoxb1b* (K), *hoxb4a* (L), *hoxb7a* (M) and *hoxb9a* (N) at 6 somite stage. Scale bar, 200 µm.

**Figure S2.**
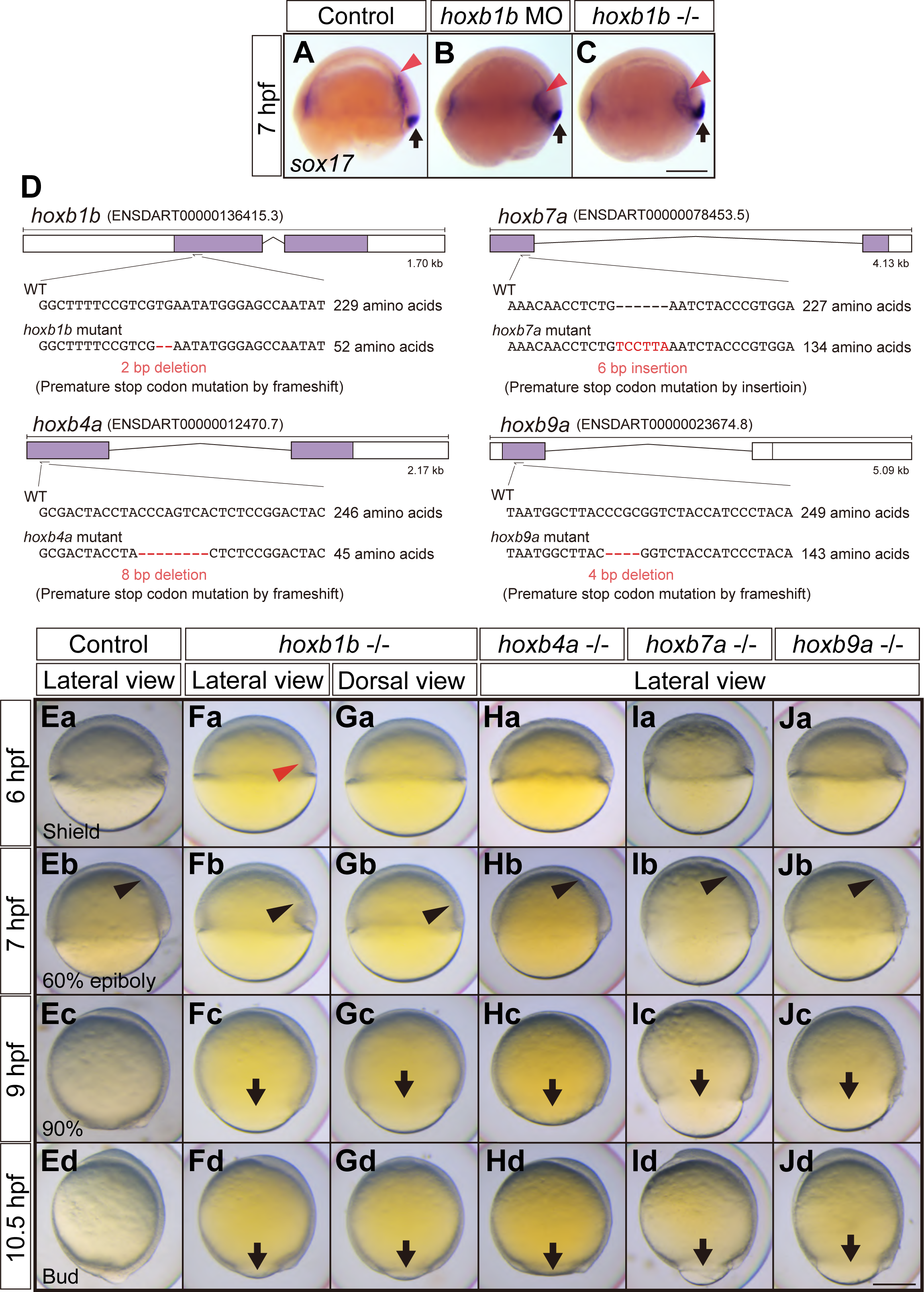
Mesendoderm formation and epiboly movements in *hoxb* mutants - Related to Figure 2. (A-C) *sox17* expression patterns in control, *hoxb1b* knock down and *hoxb1b* -/- embryos at 7 hpf. Endodermal cells migration towards the animal pole is reduced in *hoxb1b* knock down and *hoxb1b* -/- embryos. Red arrowheads point at the anterior limit of the *sox17* expression domain. Arrows depict dorsal forerunner cells. Scale bar, 200 µm. (D) Design of CRISPR/Cas9-mediated knock-out for each *hoxb* gene and resultant mutated sequences. Mutants were generated by introducing a premature stop codon through frameshift (*hoxb1b*, *hoxb4a* and *hoxb9a*) or stop codon insertion (*hoxb7a*). (E-J) Bright-field images of control (E), maternal-zygotic (MZ) *hoxb1b* (F – lateral view and G – dorsal view), MZ*hoxb4a* (H, lateral views), MZ*hoxb7a* (I, lateral views) and MZ*hoxb9a* (J, lateral views) mutants at 6 (a), 7 (b), 9 (c) and 10.5 (d) hpf. Red arrowheads point at defective mesedndoderm ingression at the dorsal blastoderm margin at 6 hpf. Black arrowheads point at the leading edge of mesendodermal cells migrating towards the animal pole. Arrows, the blastoderm margin for the embryos exhibiting epiboly delay. Dorsal side is to the right. Scale bar, 200 µm.

**Figure S3.**
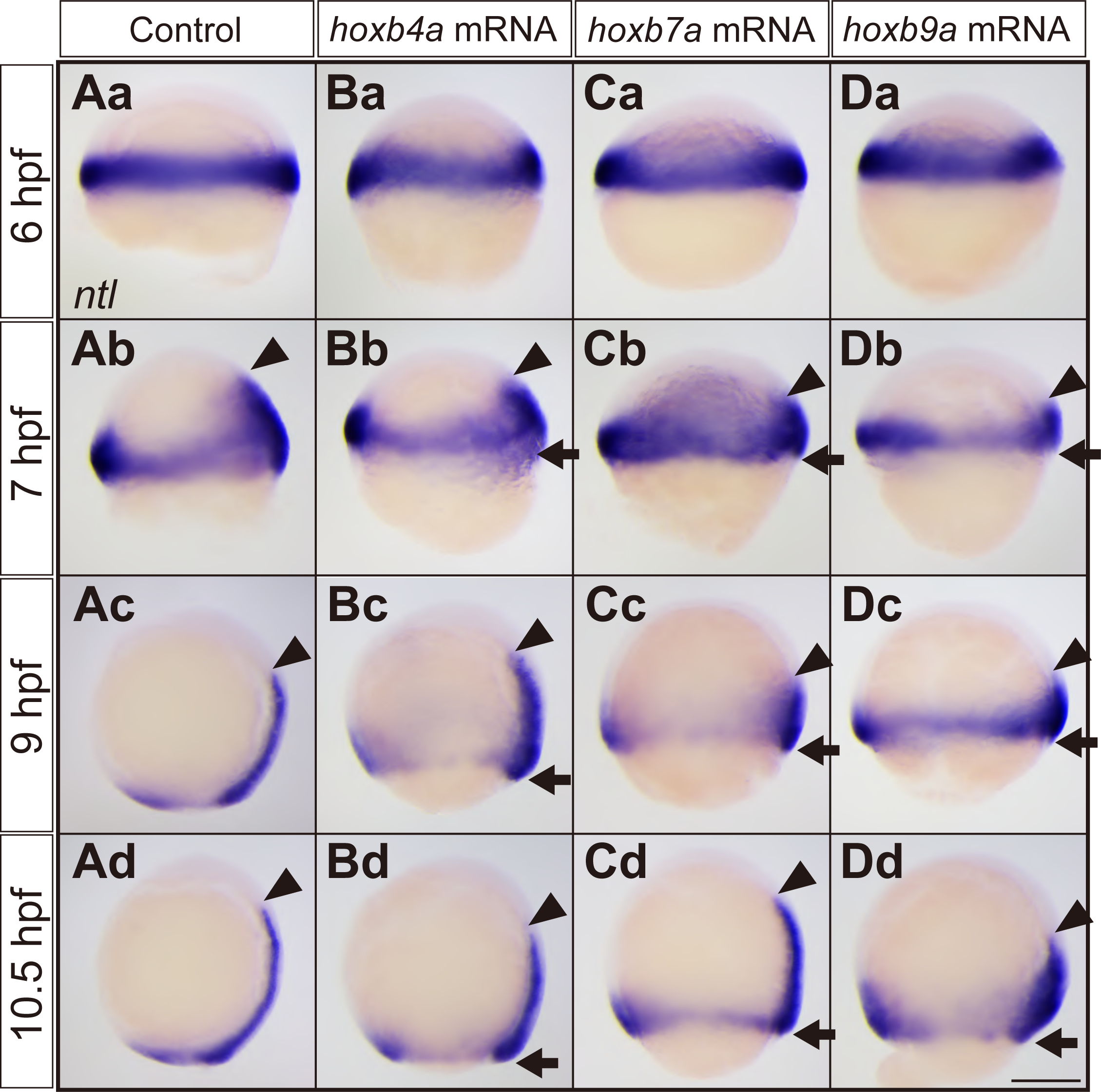
*ntl* expressions in control and *hoxb4a*, *hoxb7a* or *hoxb9a* mRNA injected embryos - Related to Figure 3. (A-D) *ntl* expressions in control (A), *hoxb4a* mRNA (B), *hoxb7a* mRNA (C) and *hoxb9a* mRNA (D) injected embryos at 6 hpf (a), 7 hpf (b), 9 hpf (c) or 10.5 hpf (d). Arrowheads demarcate the anterior limit of *ntl* expression at the dorsal side. Arrows point at the blastoderm margin in embryos with delayed epiboly. Lateral views. Dorsal side is to the right. Scale bar, 200 µm.

**Figure S4.**
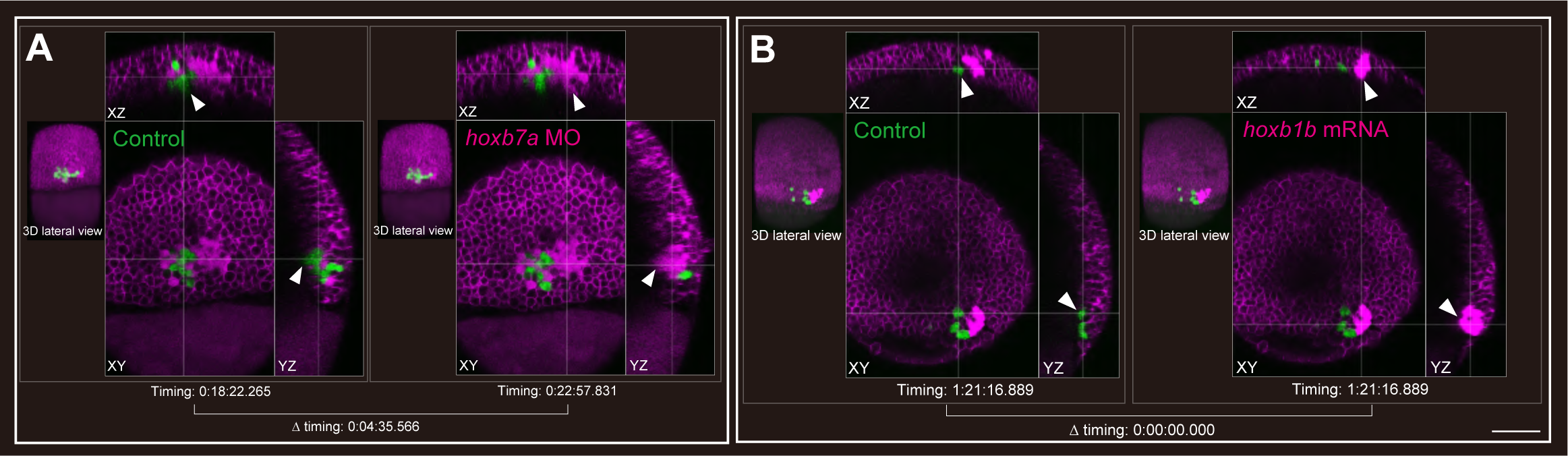
Ingression of *hoxb7a* morphant and *hoxb1b* overexpressing mesendodermal cells - Related to Figure 6. (A, B) Time needed for mesendodermal cells to complete their ingression for control cells (green, left) co-transplanted with *hoxb7a* MO (A) or *hoxb1b* mRNA (B) injected cells (magenta, right). Scale bar, 100 µm.

**Figure S5.**
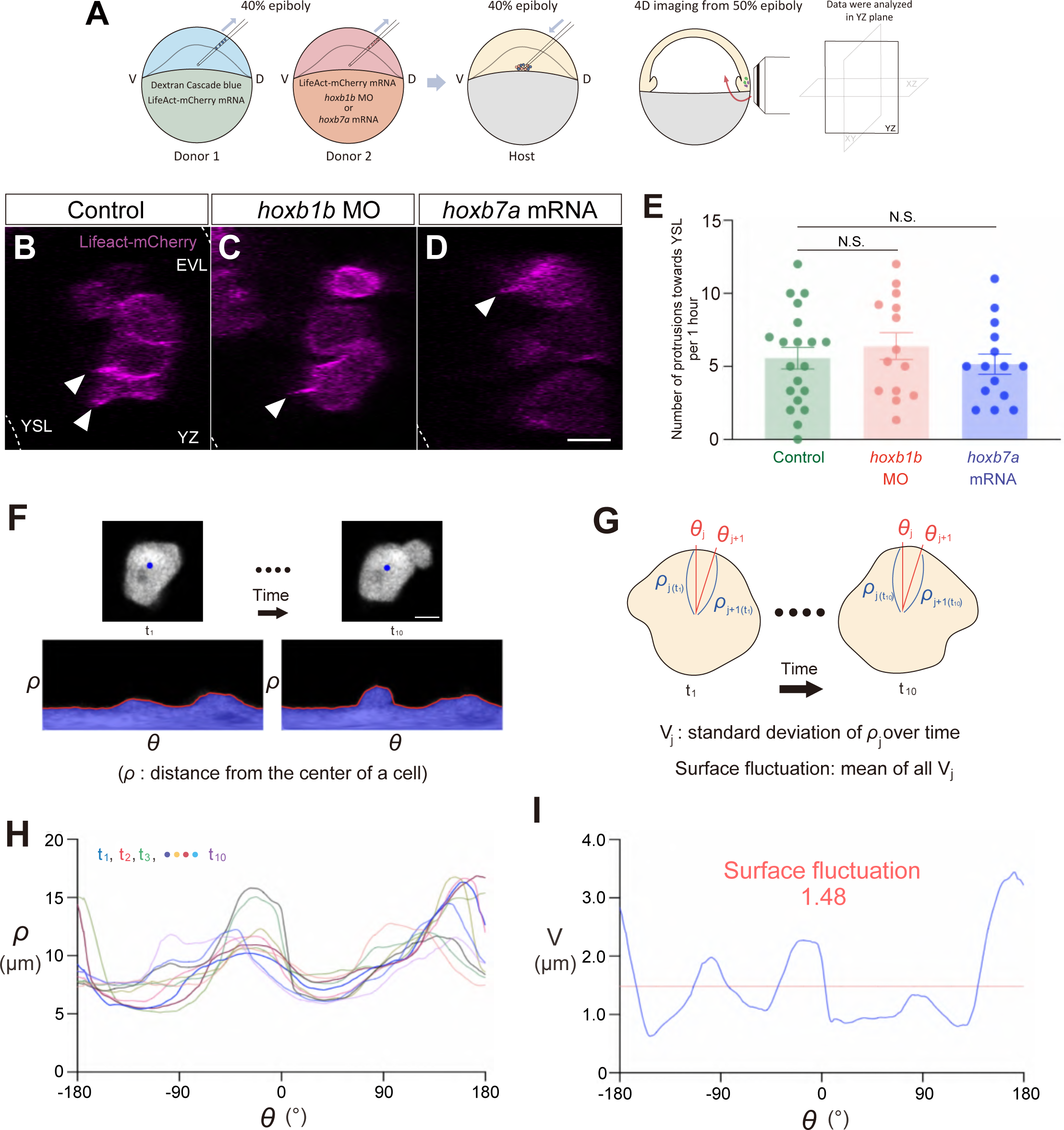
Analysis of protrusive activity and cell surface fluctuations during ingression - Related to Figure 7. (A) Schematic representation of the double transplantation assay for determining mesendodermal cell protrusive activity during ingression. Cells in the lateral blastoderm margin from dextran cascade blue and Lifeact-mCherry mRNA injected embryos (donor 1) and a combination of Lifeact-mCherry mRNA and *hoxb1b* MO or *hoxb7a* mRNA injected embryos (donor 2) were transplanted into the corresponding region of host embryos. (B-D) High-resolution confocal fluorescence images of transplanted control (B), *hoxb1b* MO (C) and *hoxb7a* mRNA (D) injected cells labelled with Lifeact-mCherry mRNA. Arrowheads point at cell protrusions directed towards the YSL (left side). Scale bar, 10 µm. YSL, yolk syncytial layer. EVL, enveloping layer. (E) Average number of protrusions directed towards the YSL formed per hour in control (n = 20, N = 6), *hoxb1b* MO (n = 12, N = 4) and *hoxb7a* mRNA (n = 15, N = 4) injected cells. Mean±s.e.m.; N.S. (not significant) with p>0.05 (*t*-test). (F-I) Quantification of cell surface fluctuations in mesendodermal cells. Fluorescence images of single cells were binarized and the distance of the margin to the center of the cell (ρ) was measured around the cell circumference (F). Surface fluctuations were calculated by determining the mean of the standard deviation of ρ_j_ over time (V_j_) (G). Representatives of calculated ρ (H) in each time points (t_1_ to t_10_) and V (I) as a standard deviation of ρ around the cell circumference and surface fluctuation as the mean of all V. Scale bar, 6 µm.

**Video S1. Control cells transplanted to the lateral germ ring margin ingressed in close succession manner - Related to Figure 6**

Time-lapse fluorescence movie for visualizing cell ingression for control (green) and control (magenta) cells transplanted to the lateral germ ring margin at 40% epiboly stage. Imaging was conducted from 50% epiboly stage (lateral view). Cell membranes of host embryo were labelled by injection of memRFP mRNA. Time-interval, 5 min 9 sec. Duration, 3 hours 21 min. Scale bar, 100 µm.

**Video S2. *hoxb1b* morphant cells transplanted to the lateral germ ring margin exhibited delayed ingression compared with these of control - Related to Figure 6**

Time-lapse fluorescence movie for visualizing cell ingression for control (green) and *hoxb1b* morphant (magenta) cells transplanted to the lateral germ ring margin at 40% epiboly stage. Imaging was conducted from 50% epiboly stage (lateral view). Cell membranes of host embryo were labelled by injection of memRFP mRNA. Time-interval, 4 min 47 sec. Duration, 3 hours 6 min. Scale bar, 100 µm.

**Video S3. *hoxb7a* overexpressing cells transplanted to the lateral germ ring margin exhibited delayed ingression compared with these of control - Related to Figure 6**

Time-lapse fluorescence movie for visualizing cell ingression for control (green) and *hoxb7a* overexpressing (magenta) cells transplanted to the lateral germ ring margin at 40% epiboly stage. Imaging was conducted from 50% epiboly stage (lateral view). Cell membranes of host embryo were labelled by injection of memRFP mRNA. Time-interval, 4 min 43 sec. Duration, 3 hours 18 min. Scale bar, 100 µm.

**Video S4. Transplanted control cells enter frequent blebbing state while co-transplanted hoxb1b morphant cells did not and exhibited static state - Related to Figure 7**

Time-lapse fluorescence movie for visualizing cell blebbing behavior for control (green) and *hoxb1b* morphant (magenta) cells transplanted to the lateral germ ring margin at 40% epiboly stage. Imaging was conducted from 50% epiboly stage (lateral view). Time-interval, 30 sec. Duration, 20 min. Scale bar, 60 µm.

**Video S5. Transplanted *hoxb1b* morphant cells delayed entering frequent blebbing state compared with these of control - Related to Figure 7**

Time-lapse fluorescence movie for visualizing cell blebbing behavior for control (green) and *hoxb1b* morphant (magenta) cells transplanted to the lateral germ ring margin at 40% epiboly stage. This movie captures the same sample as Video S4 and is a series of consecutive movie capturing 60% epiboly stage (lateral view). Time-interval, 30 sec. Duration, 20 min. Scale bar, 60 µm.

**Video S6. Control cells enter frequent blebbing state while *hoxb7a* overexpressing cells did not and exhibited static state - Related to Figure 7**

Time-lapse fluorescence movie for visualizing cell blebbing behavior for control (green) and *hoxb1b* morphant (magenta) cells transplanted to the lateral germ ring margin at 40% epiboly stage. Imaging was conducted from 50% epiboly stage (lateral view). Time-interval, 30 sec. Duration, 20 min. Scale bar, 60 µm.

**Video S7. *hoxb7a* overexpressing cells delayed entering frequent blebbing state compared with these of control - Related to Figure 7**

Time-lapse fluorescence movie for visualizing cell blebbing behavior for control (green) and *hoxb1b* morphant (magenta) cells transplanted to the lateral germ ring margin at 40% epiboly stage. This movie captures the same sample as Video S6 and is a series of consecutive movie capturing 70% epiboly stage (lateral view). Time-interval, 30 sec. Duration, 20 min. Scale bar, 60 µm.

### STAR METHODS

Detailed methods are provided in the online version of this paper and include the following:

- KEY RESOURCES TABLE
- RESOURCE AVAILABILITY

- Lead contact
- Material availability
- Data and code availability
- EXPERIMENTAL MODEL AND SUBJECT DETAILS

- Zebrafish lines
- METHOD DETAILS

- Clonings
- Whole-mount in situ hybridization
- Cryosectioning of whole-mount in situ hybridization samples
- Morpholinos and mRNAs injection
- CRISPR/Cas9 mutant generation
- Cell lineage tracing with DiI
- Visualization of intercellular spaces
- Double cell transplantation
- Imaging
- Analysis of cell movements
- QUANTIFICATION AND STATISTICAL ANALYSIS

- Statistical *analysis*

## KEY RESOURCES TABLE

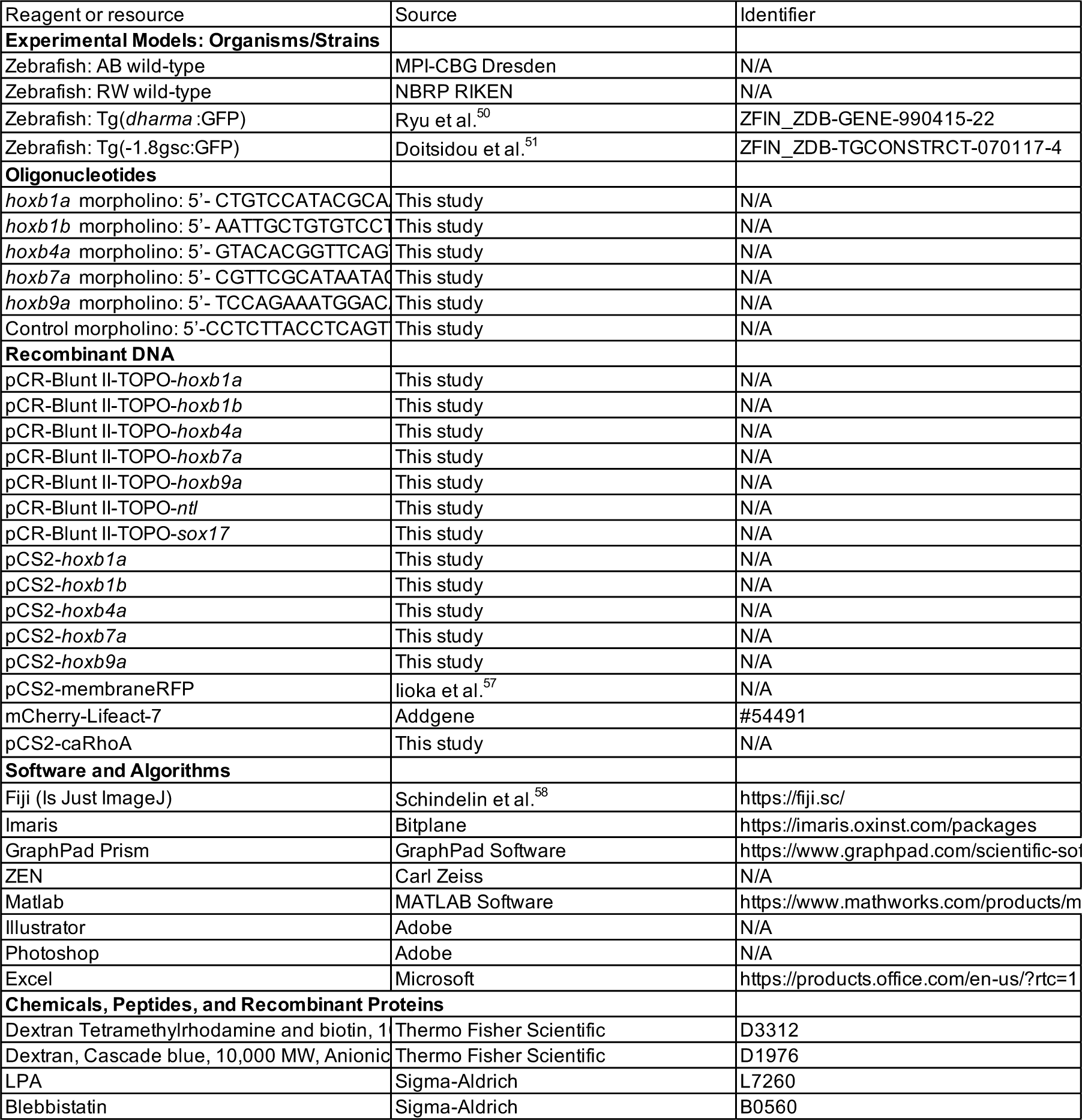

### RESOURCE AVAILABILITY

#### Lead contact

Further information and requests for resources, reagents, data, and code should be directed to the lead contact, Yuuta Moriyama.

#### Material availability

This study did not generate new unique materials nor reagents.

#### Data and code availability

No new datasets reported in this study. Code used for data analysis and any additional information in this study are available from the lead contact upon request.

### EXPERIMENTAL MODEL AND SUBJECT DETAILS

#### Zebrafish lines

Fish maintaining, injection and staging were conducted according to established procedures ^48^. AB or RW strains were used as wild-type control embryos. Embryos were obtained through natural spawning, kept in E3 medium at 28.5 °C - 31 °C prior to experiments, and staged based on morphological criteria ^49^ and hours post-fertilization (hpf). For the double transplantation assay, we used Tg(*dharma*:GFP) ^50^ or Tg(-1.8gsc:GFP) ^51^ embryos to visualize the dorsal part of the host embryos. Embryonic manipulations were conducted in Danieau’s solution [58 mM NaCl, 0.7 mM KCl, 0.4 mM MgSO_4_, 0.6 mM Ca(NO_3_) _2_, 5.0 mM HEPES (pH 7.2)] or Ringer’s solution [116 mM NaCl, 2.9 mM KCl, 1.8 mM CaCl_2_, 5.0 mM HEPES, pH 7.2] unless otherwise described. All zebrafish husbandry, breeding and experiments using zebrafish in this study were approved by the Ethics Committee of IST Austria and Aoyama Gakuin University.

### METHOD DETAILS

#### Constructs

For the cloning of *hoxb*, *ntl* and *sox17* genes for *in situ* hybridization, the following primer sets were used: *hoxb1a* (5’-CAGCACAACATCAGCACCAG - 3’, 5’-GTTGAGCTCAAGTGTGGCAG - 3’), *hoxb1b* (5’-TACTGCTTAGAGGTTGTAGG - 3’, 5’-GATAGTGGCTTGCAGAGACC - 3’), *hoxb4a* (5’-ACTCTCCGGACTACTACAGC - 3’, 5’-CAGATAGGCATAGTGTATGG - 3’), *hoxb7a* (5’-GCACCGGTCTCTTCATCATC - 3’, 5’-GTCGCCTCCAATTTGATCAG - 3’), *hoxb9a* (5’-TACCATCCCTACATACCGAC - 3’, 5’-AACGCCTAGTACCAGTCTGC - 3’). *ntl* (5’-GTCTCGACCCTAATGCAATG - 3’, 5’-CTCACAGTACGAACCCGAGG - 3’), *sox17* (5’-GCTCAGTCATGATGCCTGGC - 3’, 5’-GTAGGCAAGCTGTAGGACTC -3’). Amplified DNA fragments were cloned into pCR-Blunt II-TOPO vector using Zero Blunt TOPO PCR Cloning kit (Thermo Fisher). For the full length cloning of *hoxb* genes for mRNA injection, the following primer sets were used: *hoxb1a* (5’-TCTCGATTTCTCAGGTTGTC-3’, 5’-CGCCACCGTGTACACTGAAG-3’), *hoxb1b* (5’-ATTTATAGCTCGCTAAACAAATAC-3’, 5’-TTTATCAGGTCACAGGTCCG-3’), *hoxb4a* (5’-ATGGCCATGAGTTCCTATTTG-3’, 5’-TTGACAAAACAGAGCGACTC-3’), *hoxb7a* (5’-ATGAGTTCATTGTATTATGCGAACGCGC-3’, 5’-GTAGTTTATACATCTATATTAAAATG-3’) and *hoxb9a* (5’-AATCACTTCGCTCGAAGCGC-3’, 5’-TTTGTGGTCTATATGTAAAC-3’). Full length *hoxb* genes were subcloned into pCS2 vector. The zebrafish caRhoA plasmid was constructed as previously described ^31^.

#### Whole-mount in situ hybridization

Embryos were dechorionated using watchmaker forceps in Danieau’s or Ringer’s solution and fixed at different stages by 4% paraformaldehyde (in PBS) for 2 h at room temperature (RT) and then overnight at 4 °C. After fixation, embryos were washed three times in PBS for 10 min at RT, then transferred into methanol of increasing concentrations (25%, 50%, 75% methanol/PBS and 100% methanol) and stored at -20 °C overnight. Probes for *in situ* hybridization were synthesized from cloned cDNA sequences (described above) using SP6 RNA polymerase (10810274001, Roche) or T7 RNA polymerase (10881767001, Roche) with DIG labelling mix (11277073910, Roche). Whole-mount *in situ* hybridization for zebrafish embryos was performed as described previously ^52^ except that BM Purple (11442074001, Roche) was used for coloration. After completion of *in situ* hybridization process, samples were transferred into glycerol of increasing concentrations (25%, 50%, 75% glycerol/PBS and 100% glycerol) and then imaged on a Leica MZ165FC microscope.

#### Cryosectioning of whole-mount *in situ* hybridization samples

Stained embryos were transferred into PBS of increasing concentrations (75%, 50%, 25% glycerol/PBS and 100% PBS) and then incubated in sucrose of increasing concentrations (5%, 15%, 30% (w/v) sucrose/PBS solution) at 4 °C overnight. Samples were then placed in a 1:1 solution of 30% sucrose and optimal cutting temperature compound (OCT, Tissue-Tek) for 2 h at 4 °C, then embedded in disposal moulds with OCT and frozen at -80 °C. Frozen samples embedded in OCT were sectioned with 12 µm thickness on a Leica CM 3050 S cryostat. Sections were mounted on glass slides and allowed to dry at RT. Dried samples were washed in PBS and mounted in 75% glycerol/PBS.

#### Morpholino and mRNAs injection

Zebrafish embryos were injected using glass capillary needles (30-0020, Harvard Apparatus), which were pulled using a needle puller (P-97 IVF, Sutter Instrument) and attached to a microinjection system (PV820, World Precision Instruments). Antisense *morpholino* oligonucleotides (MOs) were designed to block translation and purchased from Gene Tools, LLC (Philomath). The sequences of MOs used in this study are as follows: *hoxb1a* (5’-CTGTCCATACGCAATTAATGGCGGA-3’), *hoxb1b* (5’-AATTGCTGTGTCCTGTTTTACCCAT-3’), *hoxb4a* (5’-GTACACGGTTCAGTATCCATATTTC-3’), *hoxb7a* (5’-CGTTCGCATAATACAATGAACTCAT-3’), *hoxb9a* (5’-TCCAGAAATGGACATTCTCGGACAT-3’) and control (5’-CCTCTTACCTCAGTTACAATTTATA-3’, supplied as standard control morpholino oligo by Gene Tools, LLC.). 2 ng of these MOs were injected into one-cell stage embryos. mRNAs of *hoxb* genes cloned in pCS2 vector were synthesized using mMESSAGE mMACHINE sp6 kit according to the manufacture’s instructions (AM1340, Invitrogen) and 100 pg of mRNA was injected into one-cell stage embryos. Lifeact-mCherry sequence was amplified from mCherry-Lifeact-7 plasmid (#54491, Addgene) with primers containing sp6 promoter sequence in the forward primer: (5’-<u>ATGACTTTAGGTGACACTATAGAAGTG</u>CTGGTTTAGTGAACCGTCAG’, 5’-AACACTCAACCCTATCTCGG-3’). Amplified DNA fragments containing sp6 promoter sequence and Lifeact-mCherry was used as template for *in vitro* transcription (mMessage mMachine Kit, Ambion). 100 pg memRFP, 50 pg Lifeact-mCherry, or 1 pg caRhoA mRNAs were injected into one-cell stage embryos.

#### CRISPR/Cas9 mutant generation

For target site determination of CRISPR/Cas9 mutants for *hoxb1b* (ENSDARG00000054033), *hoxb4a* (ENSDARG00000013533), *hoxb7a* (ENSDARG00000056030), and *hoxb9a* (ENSDARG00000056023) genes, the CHOPCHOP website was used (http://chopchop.cbu.uib.no) ^53,54^. In all cases, target sites were designed in the first exon of each gene. To generate gRNA, the following oligonucleotide sets were designed: *hoxb1b* (5’-<u>TAGG</u>ggATTGGCTCCCATATTCACGA-3’, 5’-<u>AAAC</u>TCGTGAATATGGGAGCCAATcc-3’), *hoxb4a* (5’-<u>TAGG</u>ggAGTCCGGAGAGTGACTGGGT-3’, 5’-<u>AAAC</u>ACCCAGTCACTCTCCGGACTcc-3’), *hoxb7a* (5’-<u>TAGG</u>ggTCATCCACGGGTAGATTCGG-3’, 5’-<u>AAAC</u>CCGAATCTACCCGTGGATGAcc-3’), *hoxb9a* (5’-<u>TAGG</u>ggCGCGTCTAATGGCTTACCCG-3’, 5’-<u>AAAC</u>CGGGTAAGCCATTAGACGCGcc-3’). In the oligonucleotide sequences, the adding sequences for the ligation into the Bsa I-cut pDR274 plasmid are underlined, and the adding sequences for the high efficiency for gRNA transcription *in vitro* are shown as lowercases. To anneal and generate linker DNAs using the above-mentioned oligonucleotides, 9 µl of 100 µM oligonucleotide sets each and 2 µl of 10x M buffer (TAKARA) were mixed and incubated at 72 °C for 10 min, and then the incubator was turned off and the mixture was left for 20 min to cool down. After putting on ice, 180 µl of TE buffer was added. The linker DNAs were then cloned into Bsa I-cut pDR274 plasmid and the cloned plasmids were cut by Hind III, and transcribed by MEGAshortscript T7 Transcription kit (Ambion). 1 - 2 nl of a mixture of 0.2 µl of 350 - 400 ng/µl gRNA, 2 µl of 250 - 300 ng/µl Cas9 mRNA, 0.2 µl of 350 - 400 ng/µl tyrosinase gRNA and 0.2 µl of 1M KCl was injected into one-cell stage embryos. Tyrosinase gRNA was used G0 screening ^55^. Cas9 mRNA was synthesized by mMESSAGE mMACHINE SP6 Transcription Kit according to the manufactures instructions (Ambion). For the identification of mutant fish with germline transmission, hetero-duplex mobility assay (HMA) was used ^56^. Primer sequences used for HMA are as follows: *hoxb1b* (5’-CGTTCTCACTCAAGCAGATGAC-3’, 5’-ATGATTGATAGTGGCTTGCAGA-3’), *hoxb4a* (5’-ACCCTGCGAGGAATATTCCC-3’, 5’-TGCTGGAACGAGGGGTCTTG-3’), *hoxb7a* (5’-AGAGCAGAGGGGCTACCATC-3’, 5’-GTTTTCACAGACCTGTGCTC-3’) and *hoxb9a* (5’-TCCAATGTACAGTACTCCAGCG-3’, 5’-GGTATCGAGTATCCGTTGAAGG-3’). For the identification of the mutated sequences, the amplified PCR products were cloned into the Zero Blunt TOPO PCR vector and sequenced by the sequencing service of Eurofins Genomics. Identified mutant fish were outcrossed with wild type fish to generate heterozygous lines. Homozygous mutants were obtained by incross of heterozygous fish. Maternal-zygotic mutants were obtained by crossing homozygous females with heterozygous or homozygous males. Mutant embryos were genotyped by PCR using the primers that were used for HMA.

#### Cell lineage tracing with DiI

For cell tracing, DiI (D3911, Invitrogen. 2.5 mg/ml in dimethylsulphoxide) was injected into the lateral blastoderm margin at different stages (50%, 60% or 70% epiboly). Labelled embryos were then placed into agarose moulds and imaged on a Leica MZ165FC microscope.

#### Visualization of intercellular spaces

For visualizing intercellular space (interstitial fluid), embryos were injected with 2.5 pg of Dextran Mini-Ruby (D3312, Thermo Fisher Scientific) into intercellular spaces of the lateral blastoderm margin at 40% epiboly.

#### Double cell transplantation

For double cell transplantations (cells from two donor embryos), one donor embryos was injected with 2.5 ng of Dextran Cascade-Blue (D1976, Thermo Fisher Scientific) and the other donor embryos was injected with *hoxb1b* MO/mRNA or *hoxb7a* MO/mRNA, together with 2.5 ng of Dextran Mini-Ruby (D3312, Thermo Fisher Scientific). Host embryos were injected 100 pg of membrane bound RFP (memRFP) mRNA ^57^. For the analysis of transplanted cell localization at the 12-somites stage, the two donor embryos were prepared as described above, and the host embryos remained unlabelled. For the analysis of transplanted cell protrusive activity, one donor embryo was injected with a combination of 50 pg Lifeact-mCherry and Dextran cascade-blue, and the other donor embryo was injected with a combination of 50 pg Lifeact-mCherry and *hoxb1b* MO or *hoxb7a* mRNA. For determining the bleb frequency of transplanted cells, one donor embryos was injected with 2.5 ng of Dextran Cascade-Blue and the other donor embryo was injected with *hoxb1b* MO/mRNA or *hoxb7a* MO/mRNA, together with 2.5 ng of Dextran Mini-Ruby. All of these injections were performed in one-cell stage Tg(*dharma*:GFP) or Tg(-1.8gsc:GFP) embryos, which - once injected - were incubated at 31 °C until they had reached the 30% epiboly. Embryos were then dechorionated by forceps and transferred into 2% agarose moulds within Danieau’s or Ringer’s solution; around 10-20 cells (for determining ingression timing and distribution pattern) or 5-10 cells (for observation of transplanted cell behavior) were aspirated from the lateral blastoderm margin of the donor embryo, using a bevelled borosilicate needle with a 20 µm inner diameter (Biomedical Instruments) attached to a syringe, and then transplanted into the corresponding region of the host embryo. This procedure was repeated for the second donor embryo and the order of transplantations/donor embryos was randomized to avoid artifacts due to the order of transplantations.

#### Imaging

Bright-field imaging for developing embryos and *in situ* hybridization samples was performed on a Leica microscope MZ165FC equipped with a PLANAPO 1.0x objective using LAS version 4.8 software. For imaging, developing embryos were dechorionated by forceps, and placed in 1% agarose moulds within Danieau’s solution; *in situ* hybridization samples were transferred into glycerol and mounted in 100% glycerol. For determining cell dynamics during mesendoderm cell ingression, embryos were mounted in glass bottom dishes (MATSUNAMI) within 0.7% low melting point (LMP) agarose (16520050, Invitrogen) and imaged on a FV3000 inverted confocal microscope (Olympus). For determining the timing of transplanted mesendoderm cell ingression, transplanted embryos were mounted within 0.7% LMP agarose on a mould made by 2% agarose and imaged on a TriM Scope two-photon microscope (LaVision Bio Tec) equipped with a Chameleon Ultra II laser with Chameleon Compact OPO (Coherent), a Plan-Apochromat 20x/1.0 water-immersion objective (Zeiss) and GaAsP detectors (Hamamatsu Photonics). Mesendodermal cell ingression was monitored in a 400 x 200 µm area at the blastoderm margin with a 170 µm deep z-stack in 2 µm step sizes and time intervals were 4-6 min. Images were taken with excitation wavelength of 830 nm using a Ti-Sapphire femtosecond laser system (Coherent Chameleon Ultra) and 1,100 nm optical parametric oscillator (Coherent Chameleon Compact OPO). For determining the distribution pattern of transplanted mesendodermal cells, the transplanted embryos were incubated until they had reached the 12-somites stage and then transplanted embryos were mounted within 0.7% LMP agarose on a mould made by 2% agarose and imaged. For monitoring mesendodermal cell behavior, transplanted embryos were mounted within 0.7% LMP agarose on a mould made by 2% agarose and imaged on an upright Zeiss confocal microscope LSM 900 equipped with a Plan-Apochromat 20x/1.0 water-immersion objective (Zeiss) using a ZEN software. The range of the imaged z-stack was set to capture all transplanted cells with a step size of 2-3 µm and a time interval of 30 seconds. Sectioned *in situ* hybridization samples were imaged on an Olympus BX53 microscope using cellSens software (Olympus).

#### Analysis of cell movement

Cell movement tracks were manually obtained by FLUOVIEW (Olympus). The timing of transplanted cell ingression was determined using Imaris (version 7.4, Bitplane). Protrusions (actin-rich protrusions and blebs) of transplanted cells were manually identified using ZEN (Zeiss), Fiji (NIH) and Imaris (version 7.4, Bitplane).

#### Quantification of cell surface fluctuations

Quantification of cell surface fluctuations was performed as previously described ^26^. Cell shapes, visualized on a Zeiss LSM 900 upright confocal microscope by dextran cascade blue or dextran mini-ruby staining, were determined by binarization of single z-plane images and the distance from the center of the cell to its margin (ρ) was measured. Using a custom-written MATLAB script, the outline of cell shapes was determined and converted from Cartesian into polar coordinates. Surface fluctuations as a mean of all V_j_ were normalized by the mean of radius of each cell.

### QUANTIFICATION AND STATISTICAL ANALYSIS

#### Statistical analysis

Statistical analyses of each experiment ware mentioned in the corresponding figure legends. N represents number of independent experiments and n represents number of embryos or cells. GraphPad Prism 9 and Microsoft Excel were used for statistical analysis. Statistical significances between two groups were determined by two-tailed Student’s *t*-test. Statistically significant differences are: *p < 0.05.

## Acknowledgments

We would like to thank all the Heisenberg lab members for discussions and comments on the manuscript, and the Bioimaging and zebrafish facilities of the IST Austria for continuous support. This study was funded by the Japan Society for the Promotion of Science (JSPS) Overseas Research Fellow and JST PREST (grant number: JPMJPR214B) to Y.M.

## Author contributions

Y.M. and C.-P.H. conceived of and conceptualized the project. Y.M. and C.-P.H. designed the experiments. Y.M. performed the experiments and analysed the data. T.M. performed quantification of cell surface fluctuations. Y.M. and C.-P.H. wrote the manuscript.

